# EFEMP1 contributes to light-dependent ocular growth in zebrafish

**DOI:** 10.1101/2023.12.18.572096

**Authors:** Jiaheng Xie, Bang V. Bui, Patrick T. Goodbourn, Patricia R. Jusuf

## Abstract

Myopia (short-sightedness) is the most common ocular disorder. It generally develops after over-exposure to aberrant visual environments, disrupting emmetropization mechanisms that should match eye growth with optical power. A pre-screening of strongly associated myopia-risk genes identified through human genome-wide association studies implicates *efemp1* in myopia development, but how this gene impacts ocular growth remains unclear. Here, we modify *efemp1* expression specifically in the retina of zebrafish. We found that under normal lighting, *efemp1* mutants developed axial myopia, enlarged eyes, reduced spatial vision and altered retinal function. However, under myopia-inducing dark-rearing, compared to control fish, mutants remained emmetropic and showed changes in retinal function. *Efemp1* modification changed the expression of *efemp1*, *egr1*, *tgfb1a*, *vegfab* and *rbp3* genes in the eye, and changes the inner retinal distributions of myopia-associated EFEMP1, TIMP2 and MMP2 proteins. *Efemp1* modification also impacted dark-rearing-induced responses of *vegfab* and *wnt2b* genes and above-mentioned myopia-associated proteins. Together, we provided robust evidence that light-dependent ocular growth is regulated by *efemp1*.

**Summary:** This study shows that retina-specific modification of *efemp1* expression in zebrafish results in myopic eye, and impacts responses to myopia-inducing dark-rearing in eye growth, retinal function, and myopia-associated molecular expression and distribution, implicating light-dependent regulation of *efemp1* in ocular development.

## Introduction

Myopia (short-sightedness) is now the most common visual disorder, and is predicted to impact approximately half of the world population by 2050 (Holden et al., 2016). Although considered manageable with optical correction, the development of high levels of myopia (or pathological myopia) brings with it ocular changes that promote eye diseases that cannot be easily managed (glaucoma, cataract, myopic maculopathy, etc.) (Hayashi et al., 2010; Ikuno, 2017; Marcus et al., 2011). Thus, there is an urgent need to better understand this condition.

During development, emmetropization mechanisms use visual cues to regulate eye growth, in order to match eye size with its optic power, to achieve focused vision (Rabin et al., 1981). Extended exposure to aberrant visual environments can lead to dysregulation of emmetropization mechanisms resulting in myopia. Human studies showed that children who spent more time outdoors were less likely to develop myopia (Rose et al., 2008). Recent shifts away from outdoor activities and increased screen time may further increase myopia prevalence and promote pathological myopia (Najafzadeh et al., 2023). This points to an urgent need to understand the underlying mechanisms for myopia development, to underpin the development of novel approaches to target these mechanisms.

A range of myopia-associated genes have been identified over the past several decades (Tedja et al., 2019). Meta-analyses of an increasingly large pool of human genome-wide association (GWAS) data have seen a growth in the number of novel risk genes associated with refractive error (Tedja et al., 2018). Amongst the highest ranked myopia risk genes is *EGF containing fibulin extracellular matrix protein 1* (*efemp1*). EFEMP1 is a secreted extracellular matrix glycoprotein widely expressed throughout the human body, especially in elastic fiber-rich tissues, for examples, the brain, lung, kidney and eye including the retina (Livingstone et al., 2020; Mackay et al., 2015). Our previous study (unpublished) screening GWAS-associated myopia-risk genes with high-throughput optomotor response measurement and morpholino gene knockdown indicated that knockdown of *efemp1* in larval zebrafish reduced spatial-frequency tuning function, making *efemp1* a candidate gene worth for further investigation for myopia development. In relation to human visual diseases, an autosomal dominant gain-of-function mutation in the *EFEMP1* gene (c.1033C>T, p.Arg345Trp) is known to be associated with Malattia Leventinese and Doyne honeycomb retinal dystrophy (Stone et al., 1999), leading to accumulation of drusen (yellow-white deposits) beneath the basal retinal pigment epithelium (RPE), a pathologic sign that overlaps with age-related macular degeneration (AMD) (Marmorstein et al., 2002). A number of studies suggest that sub-RPE deposits may result from abnormal interactions between EFEMP1 and complement system proteins, for example, complement component 3 (C3), complement factor B (FB) and complement factor H (CFH) (Crowley et al., 2022; Garland et al., 2013).

Recently, from 3 independent Filipino families, three novel *EFEMP1* variants (c.238A>T, p.Asn80Tyr; c.1480T>C, p.Ter494Glnext*29; and c.1429C>T, p.Arg477Cys) were identified in patients with juvenile-onset open-angle glaucoma, in which 76% of the carriers eventually developed blindness (Collantes et al., 2022). This study also demonstrated that protein aggregation and retention was the potential cause of juvenile-onset open-angle glaucoma. In addition, a recent reported case with biallelic *EFEMP1* loss-of-function variants (c.320_324del, p.Met107fs; c.615T>A, p.Tyr205Ter) manifested connective tissue abnormalities, including tall stature, hernia, hypermobile joints, thin translucent skin, and importantly high myopia (Driver et al., 2020). These clinical findings highlight a role for *EFEMP1* in ocular development, however how *efemp1* is involved in ocular refractive development remains largely unknown.

In the current study, using a zebrafish model with retina-specific *efemp1* modification, we show robust evidence that *efemp1* regulates ocular growth in a light-dependent manner in zebrafish. In particular, mutant fish developed myopia and enlarged eyes when reared under normal lighting, but became emmetropic after dark-rearing (a myopia-inducing condition for normal fish). In addition, we show that *efemp1* may mediate ocular growth by regulating expression of *early growth response 1* (*egr1*), *transforming growth factor beta 1b* (*tgfb1a*), *vascular endothelial growth factor Ab* (*vegfab*) and *retinol binding protein 3* (*rbp3*), and modulating light-dependent responses of *Wnt family 2B* (*wnt2b*) and *vegfab*. Meanwhile, retinal *efemp1* modification influences inner retinal distribution of matrix metalloproteinase 2 (MMP2) and tissue inhibitor of metalloproteinase 2 (TIMP2) for both intrinsic and visually induced ocular growth. This study demonstrates the capacity of a comprehensive zebrafish analysis platform, combining genetic and environmental manipulations, as well as multifaceted phenotypic assessments, to gain insights into GWAS-associated myopia-risk genes in myopia development.

## Results

### Retina-specific disruption of *efemp1* leads to reduced visual sensitivity

In order to investigate the role of the *efemp1* gene and its interaction with visual environment, we first generated a zebrafish line with *efemp1* modification specifically in the retina (*efemp1*^2C-Cas9^; Fig 1A), the light-sensing tissue in the eye, using a 2C-Cas9 somatic CRISPR gene editing system (Di Donato et al., 2016). In this transgenic zebrafish line, Tg(*rx2*:*Gal4*) is expressed specifically in the retina and the RPE (Chuang and Raymond, 2001), due to the retina-specific *retinal homeobox gene 2* (*rx2*) promoter. The Gal4 transcription factor binds the upstream activating sequence (UAS) element of the 2C-Cas9 transgene Tg(*UAS*:*Cas9T2ACre*;*U6*:*efemp1sgRNA1*;*U6*:*efemp1sgRNA2*), leading to expression of the Cas9 nuclease and Cre recombinase. The retina-specific Cas9 nuclease then combines with universally expressed *efemp1* small guide RNAs (sgRNAs), which target exons 3 and 5 of the *efemp1* genome DNA (gDNA) for CRISPR gene editing in the retina (Fig 1B). The Cre recombinase recombines loxP sites in the Tg(*bact2*-*loxP*-*mCherry*-*loxP*-*eGFP*) and switches mCherry to eGFP fluorophore reporter specifically in the eye, thus allowing for selection of Cas9-positive mutants (Fig 1A′). The patchiness of Cas9 expression in the mutant retina may attribute to the Gal4/UAS system (Halpern et al., 2008). This gene editing system led to mosaic retinal mutations; each Cas9-expressing retinal cell that were driven by the *rx2* promoter would perform its own CRISPR gene editing process, and as a result, even in an individual retina, there were different types of indels (e.g., loss- or gain-of-function mutations, milder mutations that may cause mislocalization) in different cells. Despite the mosaicism, the mutations resulted from the 2C-Cas9 system in retinal cells is expected to be sustained. Also, in adult teleost, activation of *rx2* in retinal stem cells in the retinal ciliary marginal zone determines its fate to form retinal neurons (Reinhardt et al., 2015). This suggested that in new neurons derived from retinal stem cells in the adult zebrafish retina, there is expression of *rx2* to drive the 2C-Cas9 system for genetic modification. Genetic control fish (*efemp1*^+/+^) had the same transgenic background except that they lacked the 2C-Cas9 transgene.

**Figure 1.**
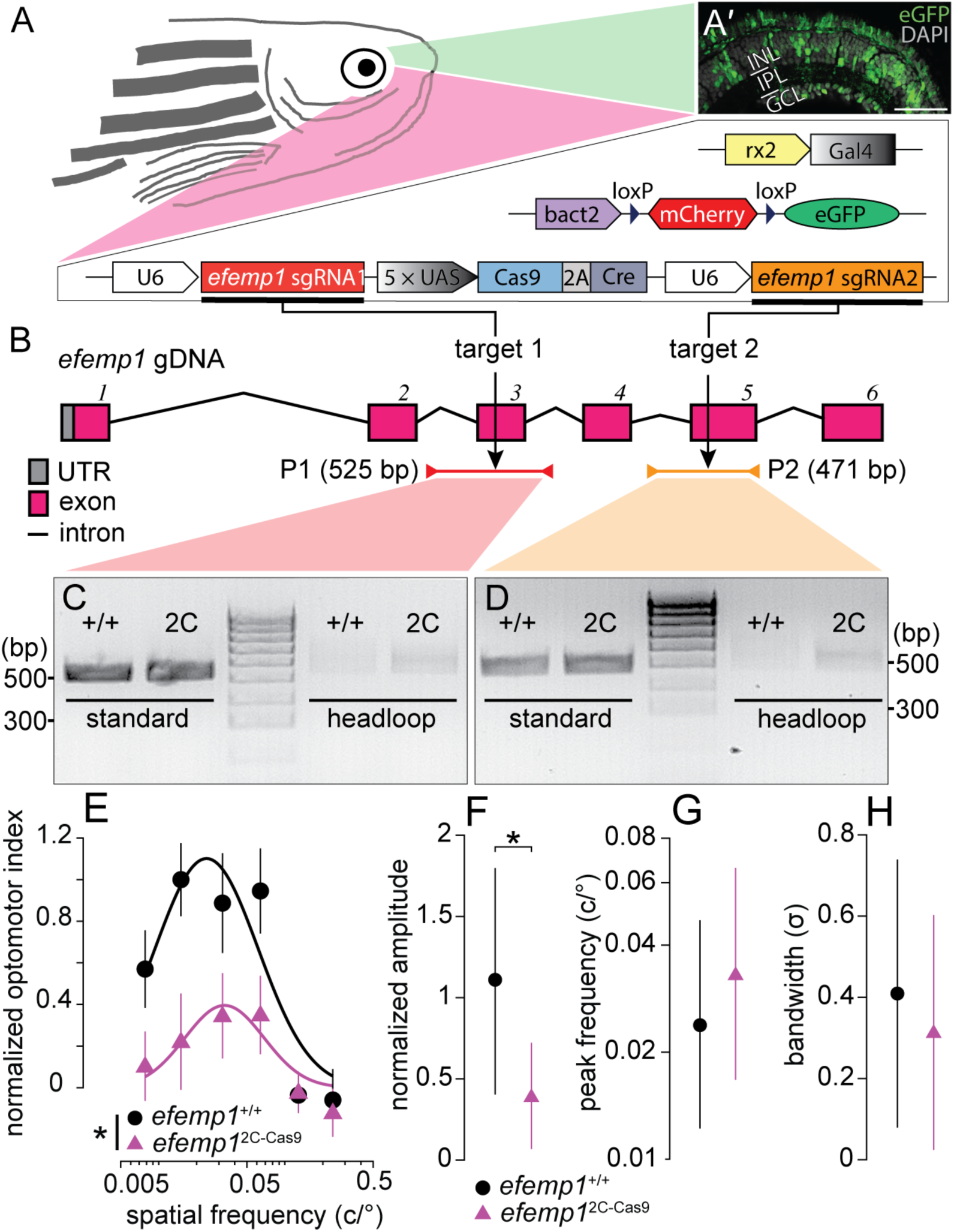
Genotypic details of retina-specific *efemp1*-modified mutants (*efemp1*^2C-Cas9^) and phenotypic verification via optomotor responses at 5 days post-fertilization (dpf). **(A)** The *efemp1*^2C-Cas9^ fish were generated using 2C-Cas9 somatic gene editing. Mutant fish have three separate transgenes: Tg(*rx2*:*Gal4*) × Tg(*bact2*-*loxP*-*mCherry*-*loxP*-*eGFP*) × Tg(*UAS*:*Cas9T2ACre*;*U6*:*efemp1sgRNA1*;*U6*:*efemp1sgRNA2*). Co-expression of these transgenic elements result in expression of green fluorescence (eGFP) in zebrafish retina, indicative of successful Cas9 protein expression. **(Aʹ)** Representative retinal image from a 6 dpf fish. Nuclei stained with DAPI are shown in grey. INL, inner nuclear layer; IPL, inner plexiform layer; GCL, ganglion cell layer. Scale bar: 40 μm. **(B)** The universally expressed *efemp1* sgRNA1 and sgRNA2 from the transgene binds with Cas9 nucleases and cuts exons 3 and 5 of the *efemp1* genome DNA (gDNA), respectively. Standard PCR primers used for genotyping the target sites 1 and 2 amplified 525- (Product 1, P1; spinning from the introns before to after the exon 3) and 471-base pairs (bp; P2; spinning from the intron before to the end of exon 5) of DNA sequences, respectively. Products of headloop PCR are 23-bp longer for P1 and 20-bp longer for P2 due to the headloop tags in reverse primers. UTR, untranslated region. **(C–D)** Gel electrophoresis of the products from standard and headloop PCR for genotyping targeted sites 1 **(C)** and 2 **(D)**. In both images, the middle lanes show a 100-bp reference ladder; positions of 300 and 500 bp of size are indicated for both gel images. **(E–H)** Results of optomotor responses. Spatial-frequency tuning functions **(E)** for 5 dpf *efemp1*^+/+^ (n = 14) and *efemp1*^2C-Cas9^ fish (n = 13) are three-parameter log-Gaussian functions fit to the data by minimizing the least-square error. The fitted parameters, including **(F)** normalized amplitude, **(G)** peak frequency and **(H)** bandwidth, were compared between groups Group data are shown as mean ± SEM in (E) and mean with 95% confidence intervals in (F–H). \**P* < *0.05* (*F*-test).

We used headloop PCR, which specifically suppresses amplification of wildtype (unmutated) DNA sequences containing sgRNA target sites, but allows targeted mutant sequences to undergo normal PCR amplification. We showed that there were discernibly higher amounts of PCR products amplified from 30 ng of *efemp1*^2C-Cas9^ eye gDNA than from 30 ng of control eye gDNA for both target sites, demonstrating that there were mutations in the mutant zebrafish retina (Fig 1C–D). However, the headloop PCR bands of *efemp1*^2C-cas9^ fish were remarkably weaker than standard PCR bands, suggesting a low mutation rate. This may have arisen as our gDNA was extracted from whole zebrafish eyes rather than just the retina or Cas9+ cells. On the other hand, due to the mosaic nature of the gene editing, the patchiness of Cas9-expressing retinal cells (Fig 1A′) and the potentially low editing rate, as well as the unavailability of commercial anti-EFEMP1 antibodies that targets specifically the CRISPR editing sites, *efemp1* modification in our mutant model at the protein level is challenging to show.

Our preliminary data showed that 5 days post-fertilization (dpf) fish with *efemp1* knockdown by morpholino presented reduced spatial-frequency tuning functions (unpublished). Therefore, we tested optomotor responses (OMR) for our mutant fish to verify modification of *efemp1* at the behavioral level. Our 5 dpf *efemp1*^2C-Cas9^ mutants showed altered spatial-frequency tuning functions (*P* = *0.0334*; Fig 1E), with a significant amplitude reduction (*P* = *0.0412*; Fig 1F) in *efemp1*^2C-Cas9^ fish. There was no difference in peak spatial frequency (Fig 1G) and function bandwidth (Fig 1H) between genotypes. OMR assessment at 2 weeks post-fertilization (wpf) returned similar differences between mutants and their wildtype counterparts (spatial-frequency tuning function: *P* = *0.0034*; amplitude: *P* = *0.0022*; Fig S1). Taken together, *efemp1*^2C-Cas9^ fish are a robust model of retinal *efemp1* modification.

### Disruption of *efemp1* causes myopia and abnormal ocular development in zebrafish

We investigated the development of ocular refraction in *efemp1*^2C-Cas9^ mutant zebrafish using optical coherence tomography (OCT) to non-invasively obtain cross-section images of zebrafish eyes. This allowed us to quantify ocular refraction as the ratio of retinal radius to lens radius (R/L ratio; Fig 2A), namely *Matthiessen’s Ratio*, which is a common means to return axial ocular refraction for aquatic species (Collery et al., 2014; Shand et al., 1999; Turnbull et al., 2015). A higher R/L ratio compared to control fish indicates relative axial myopia, while a lower ratio represents a relative hyperopic shift. Overall, *efemp1*^2C-Cas9^ fish have higher R/L ratio than *efemp1*^+/+^ fish (*P* = *0.0002*), indicating relative myopic ocular structure*. Post-hoc* analysis identified significant relative axial myopia at 2, 6 and 8 weeks of age (*P* = *0.018*, *0.0001* and *0.002*, respectively; Fig 2B).

**Figure 2.**
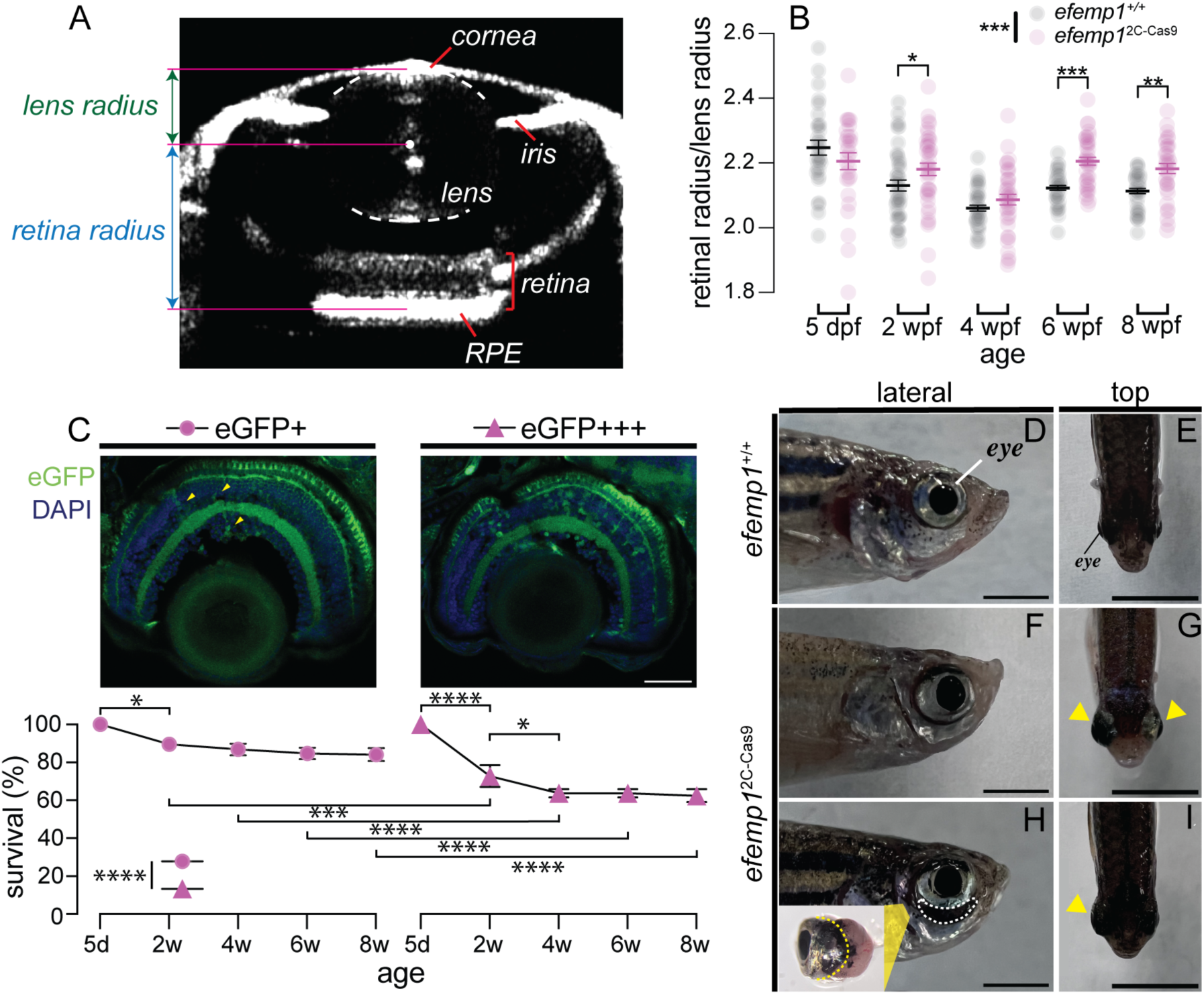
Ocular development of *efemp1*^2C-Cas9^ fish under normal rearing. **(A)** Representative optical coherence tomography (OCT) image showing eye components, including the cornea, iris, lens, retina and retinal pigment epithelium (RPE). Relative ocular refraction was calculated as the ratio of retinal radius to lens radius. **(B)** Ocular refraction of *efemp1*^+/+^ and *efemp1*^2C-Cas9^ fish at 5 days post-fertilization (dpf; n = 30 and 29 eyes, respectively), and 2 (n = 40 per genotype), 4 (n = 40 and 38 eyes, respectively), 6 (n = 39 and 40 eyes, respectively), and 8 weeks post-fertilizations (wpf; n = 40 and 34 eyes, respectively). Group data are shown as mean ± SEM. **(C)** *Efemp1*^2C-Cas9^ fish were categorized into two groups based on their retinal eGFP levels (eGFP+ vs. eGFP+++). Sporadic eGFP positive cells (highlighted by yellow arrows) are evident in the eGFP+ retinal image, whereas the eGFP+++ retinal image had more EGFP cells. Scale bar: 40 μm. Survival rates for eGFP+ and eGFP+++ groups from 5 days (d; set as 100%) to 8 weeks (w) of age are presented as mean ± SEM. Three tanks per group were analyzed. \**P* < *0.05*; \*\**P* < *0.01*; \*\*\*\**P* < *0.0001*. *Two*-way ANOVA and Fisher’s LSD *post-hoc* tests were performed. In **(C)**, for comparisons within each eGFP level, only significant survival differences between a time point and its adjacent time are shown. **(D–I)** Representative images of eye morphology for *efemp1*^+/+^ **(D–E)** and *efemp1*^2C-Cas9^ fish **(F–I)** reared under normal lighting at 1 year old. Yellow arrowheads indicate enlarged eyes. Dashed region below the eye in **(H)** highlights a scleral crack shown in the zoom-in image insert, in which the yellow dashed line indicates the estimated normal position of the posterior eye of zebrafish. Scale bars: 3 mm.

We considered why axial myopia was not significant at 4 wpf in *efemp1*^2C-Cas9^ fish. EGFP fluorescence levels in the eye appeared to vary between *efemp1*^2C-Cas9^ fish. We speculated that *efemp1*^2C-Cas9^ fish with higher eGFP expression indicating higher Cas9 nuclease expression and thus higher rate of CRISPR gene editing, would have more severe visual impairment. As visual impairment can impact feeding, we examined survival rate in low (eGFP+) and high eGFP (eGFP+++) expressing *efemp1*^2C-Cas9^ fish, differences that were easily distinguished via *post-hoc* histology (Fig 2C). Those *efemp1*^2C-Cas9^ fish with weaker retinal eGFP expression had higher survival rates than GFP+++ group (*P* < *0.0001*; Fig 2C). About 10% of GFP+ fish were lost by 2 wpf (*P* = *0.020*) but survival was stable thereafter. In contrast, GFP+++ fish had significantly lower survival rates than GFP+ group at 2 and 4 wpf (*P* = *0.0007* and *P* < *0.0001*, respectively). The survival rate stabilized in GFP+++ from 4 wpf. We interpret these data to indicate that at 4 wpf, loss of the more myopic fish may have contributed to the absence of a significant effect at this age (Fig 2C).

We also found that at 5–8 months of age, 7.11% of *efemp1*^2C-Cas9^ fish appear to have monocular or binocular enlargement (154 fish in total from 5 tanks; Table S1; Fig 2F–I, yellow arrows), compared to *efemp1*^+/+^ fish (Fig 1D–E). In some cases, there was a dark crescent between the eye socket and the ventral eyeball (Fig 2H, white dashed area). Dissection of this eye showed aggressive posterior eye expansion and a scleral crack (Fig 2H insert). Further study is required to confirm whether these rare, severe scleral cracks represented a greater or faster ocular expansion caused by our retinal *efemp1* modification. Overall, the above results suggest that disruption of *efemp1*^2C-Cas9^ leads to myopic vision, eye enlargement, and possibly more severe ocular pathology.

### EFEMP1 deficits in the retina results in altered retinal function

The retina plays a crucial role in the local regulation of eye growth, a process that does not require input from the brain (Troilo et al., 1987; Wallman et al., 1987). To examine whether retinal function is involved in the abnormal development of ocular refraction in the *efemp1*^2C-Cas9^ fish, electroretinograms (ERG) were recorded at 2, 4, 6 and 8 wpf.

At 2 weeks of age, *efemp1*^2C-Cas9^ fish showed significantly lower photoreceptoral *a*-wave amplitude (*P* = *0.0118*), especially at higher stimulus intensities (Fig 3A_2_), compared to *efemp1*^+/+^ fish. However, *b*-wave amplitude was similar between mutants and control fish (Fig 3A_1_). *A*- and *b*-wave implicit times of *efemp1*^2C-Cas9^ fish were significantly faster than that of *efemp1*^+/+^ fish (*P* = *0.018* and *0.0047*, respectively; Fig 3A, 4A_3_–A_4_).

**Figure 3.**
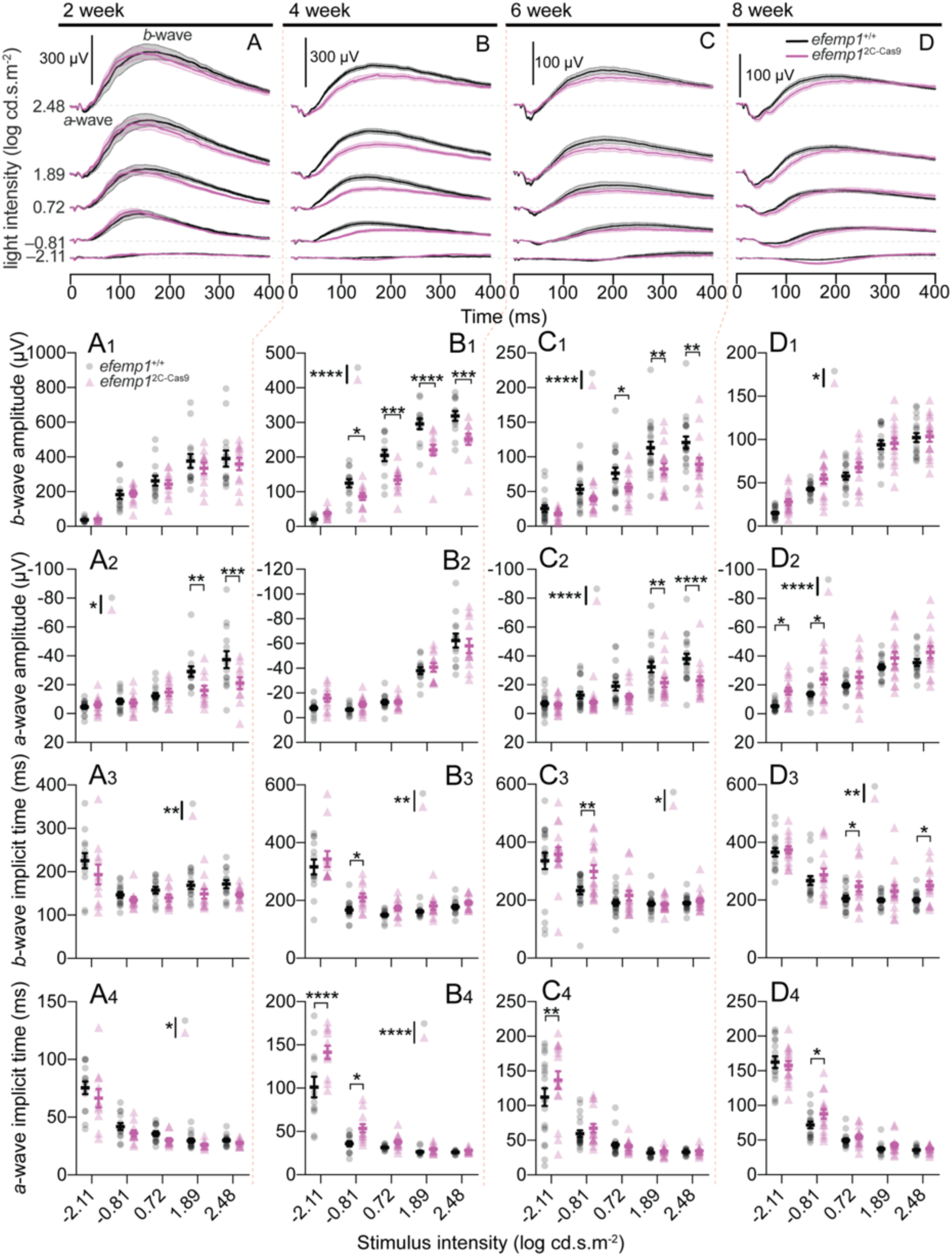
Electroretinogram (ERG) results of *efemp1*^+/+^ and *efemp1*^2C-Cas9^ fish under normal rearing. **(A, B, C, D)** Group average ERG traces with shades areas indicating SEM, **(A_1_, B_1_, C_1_, D_1_)** *b*-wave amplitudes, **(A_2_, B_2_, C_2_, D_2_)** *a*-wave amplitudes, **(A_3_, B_3_, C_3_, D_3_)** *b*-wave implicit times and **(A_4_, B_4_, C_4_, D_4_)** *a*-wave implicit times at 2 **(A–A_4_, respectively)**, 4 **(B–B_4_, respectively)**, 6 **(C–C_4_, respectively)** and 8 **(D–D_4_, respectively)** weeks post-fertilization (wpf). There were 14 and 11 eyes for 2 wpf, 13 and 12 eyes for 4 wpf, 20 and 17 eyes for 6 wpf, and 17 and 16 eyes for 8 wpf *efemp1*^+/+^ and *efemp1*^2C-Cas9^ fish, respectively. Scale bars: 300 µV in **(A–B)** and 100 µV in **(C–D)**. Group data are shown as mean ± SEM. *T*wo-way ANOVA and Fisher’s LSD *post-hoc* tests were performed. \**P* < *0.05*; \*\**P*< *0.01*; \*\*\**P* < *0.001*; \*\*\*\**P* < *0.0001*.

At 4 wpf (Fig 3B–B_4_), *efemp1*^2C-Cas9^ fish showed smaller *b*-wave amplitude (*P* < *0.0001*; Fig 3B_1_), but no difference in *a*-wave amplitude (Fig 3B_2_) compared with *efemp1*^+/+^ fish. *Two*-way ANOVA also indicated significantly slower *b*-wave (*P* = *0.0052*, Fig 3B_3_) and *a*-wave (*P* < *0.0001*, Fig 3B_4_) in *efemp1*^2C-Cas9^ fish.

At 6 wpf, both *a*- and *b*-wave amplitudes were significantly smaller in *efemp1*^2C-Cas9^ relative to *efemp1*^+/+^ fish (effects of genotypes: *P* < *0.0001* for both *a*- and *b*-wave; Fig 3C–C_2_). *Efemp1*^2C-Cas9^ fish had significantly slower *b*-wave (effect of genotypes: *P* = *0.0182,* Fig 3C_3_) and *a*-wave responses (*P* = *0.0068* for –2.11 log cd.s.m^-2^; Fig 3C_4_) than *efemp1*^+/+^ fish.

At 8 wpf (Fig 3D), both *b*-wave (*P* = *0.012*; Fig 3D_1_) and *a*-wave amplitudes (*P* < *0.0001*; Fig 3D_2_) were a little larger in *efemp1*^2C-Cas9^ compared to *efemp1*^+/+^ fish. A clear effect of genotypes was evident in terms of slower *b*-waves (*P* = *0.0012*; Fig 3D_3_) and *a*-waves (*P* = *0.015*; Fig 3D_4_) in *efemp1*^2C-Cas9^ fish. A longer latency before initiation of the *b*-wave might expose a larger proportion of the electro-negative photoreceptoral *a-wave* responses, accounting for the observed larger but slower *a-waves*.

Although it is unclear why *efemp1*^2C-Cas9^ mutants show faster response at a young age, but slower response at 4-8 wpf, it is apparent that retinal dysfunction is a robust phenotype of this genetic myopia mutant model.

### Zebrafish with retinal *efemp1* disruption become emmetropic after dark-rearing

As our results showed that *efemp1* disruption led to myopia development in zebrafish under normal lighting, we further investigated whether *efemp1* modification might exacerbate the myopia-inducing effects of dark-rearing (Fig 4A) (Xie et al., 2023 Preprint).

**Figure 4.**
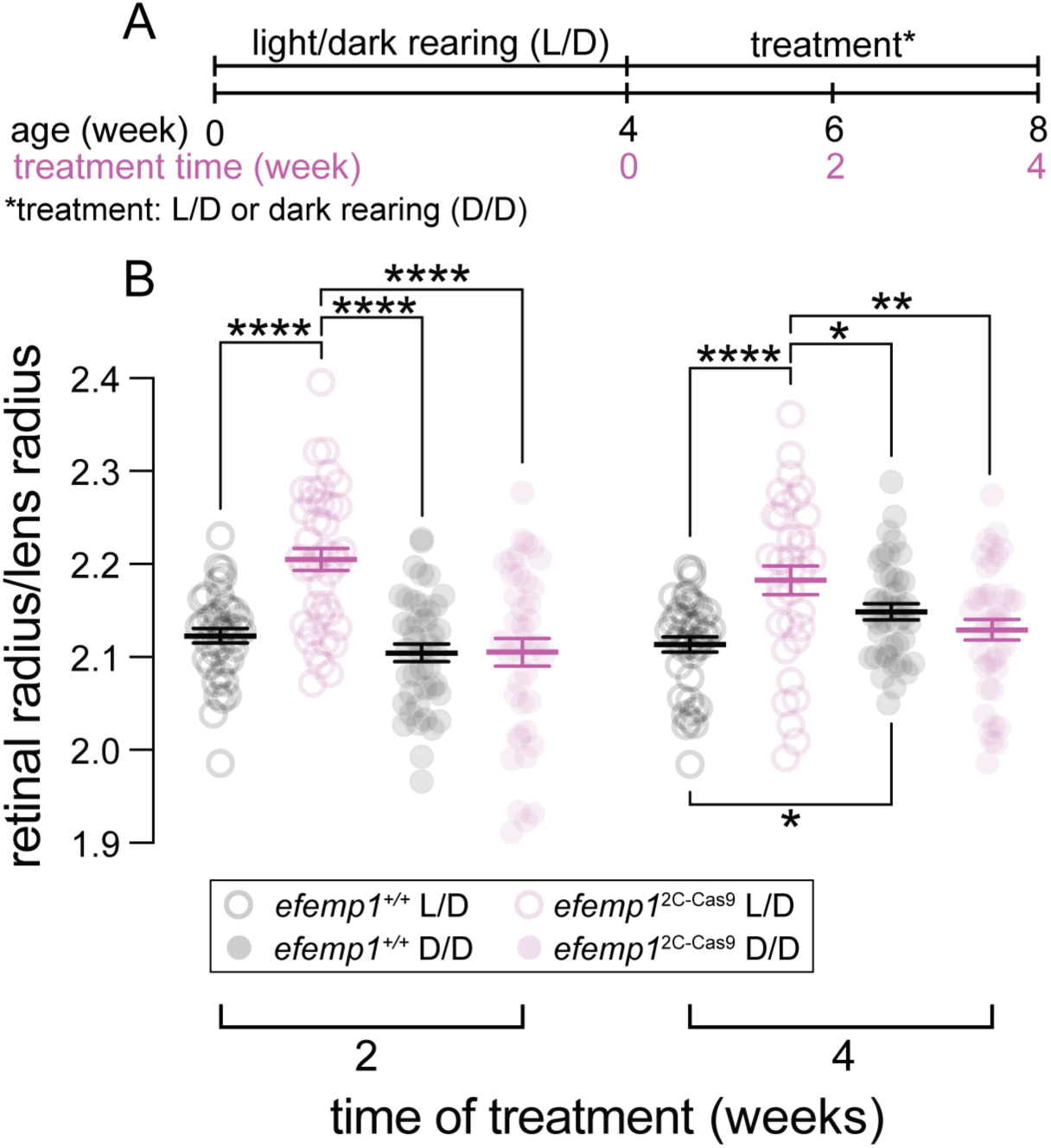
Ocular refraction of *efemp1*^+/+^ and *efemp1*^2C-Cas9^ fish after environmental treatment. **(A)** *Efemp1*^+/+^ and *efemp1*^2C-Cas9^ fish were reared under normal light/dark cycle (L/D) until 4 weeks post-fertilization (wpf) and either continued under normal lighting (L/D) or switched to dark-rearing (D/D) for 2 or 4 weeks (to 6 or 8 wpf, respectively). **(B)** Relative ocular refraction of L/D- or D/D-reared *efemp1*^+/+^ and *efemp1*^2C-Cas9^ eyes were quantified after 2 or 4 weeks of treatment using optical coherence tomography (OCT). There were 39, 40, 42 and 40 eyes at the 2-week time point, and 40, 34, 40 and 40 eyes at the 4-week time point for L/D-reared *efemp1*^+/+^, L/D-reared *efemp1*^2C-Cas9^, D/D-reared *efemp1*^+/+^, D/D-reared *efemp1*^2C-Cas9^ fish, respectively. Group data are shown as mean ± SEM. *Three*-way ANOVA and Fisher’s LSD *post-hoc* analyses were performed. \**P* < *0.05*; \*\**P* < *0.01*; \*\*\**P* < *0.001*; \*\*\*\**P* < *0.0001*.

After 2 weeks of rearing under normal lighting, *efemp1*^2C-Cas9^ fish had developed myopia compared with *efemp1*^+/+^ fish (*P* < *0.0001*). Interestingly, after 2 weeks of dark-rearing*, efemp1*^2C-Cas9^ fish did not undergo a myopic shift (*P* < *0.0001*, Fig 4B), and had similar ocular fraction to *efemp1*^+/+^ fish reared under normal light.

After 4 weeks of dark-rearing, control *efemp1*^+/+^ fish became myopic (*P* = *0.025*; Fig 4B). However, similar to 2-week results, relative to normal-light-reared *efemp1*^+/+^ fish, *efemp1*^2C-Cas9^ fish were myopic when reared under normal lighting (*P* < *0.0001*), but after dark-rearing they maintained a R/L ratio that was consistent with emmetropic vision at this age (Fig 4B).

A *three*-way ANOVA reveals a significant interaction between genotypes and environmental conditions (*P* < *0.0001*). These results indicated that under normal visual conditions *efemp1* is needed for emmetropization, however under aberrant visual conditions the lack of *efemp1* can interfere with mechanisms that usually drive myopia development.

### Retinal *efemp1* modification drives retinal physiological changes in response to dark-rearing

To investigate whether *efemp1* is involved in dark-rearing-induced retinal functional changes, we analyzed ERG of *efemp1*^+/+^ and *efemp1*^2C-Cas9^ fish after 2 days, 2 or 4 weeks of normal light or dark-rearing.

After 2 days of treatment, under either rearing condition, *efemp1*^2C-Cas9^ fish showed smaller ERG responses than *efemp1*^+/+^ fish (Fig 5A–D). *Three*-way ANOVA confirmed a significant genotype effect with smaller *b*- and *a*-wave amplitudes, as well as slowed *b*- and *a*-wave implicit times (*P* < *0.0001*, *P* = *0.0002*, *P* = *0.0023* and *P* = *0.021*, respectively; Fig 5E–H) for *efemp1*^2C-Cas9^ fish. There was a significant interaction between genotypes and rearing conditions for *b*-wave implicit time (*P* = *0.0362*; Fig 5G).

**Figure 5.**
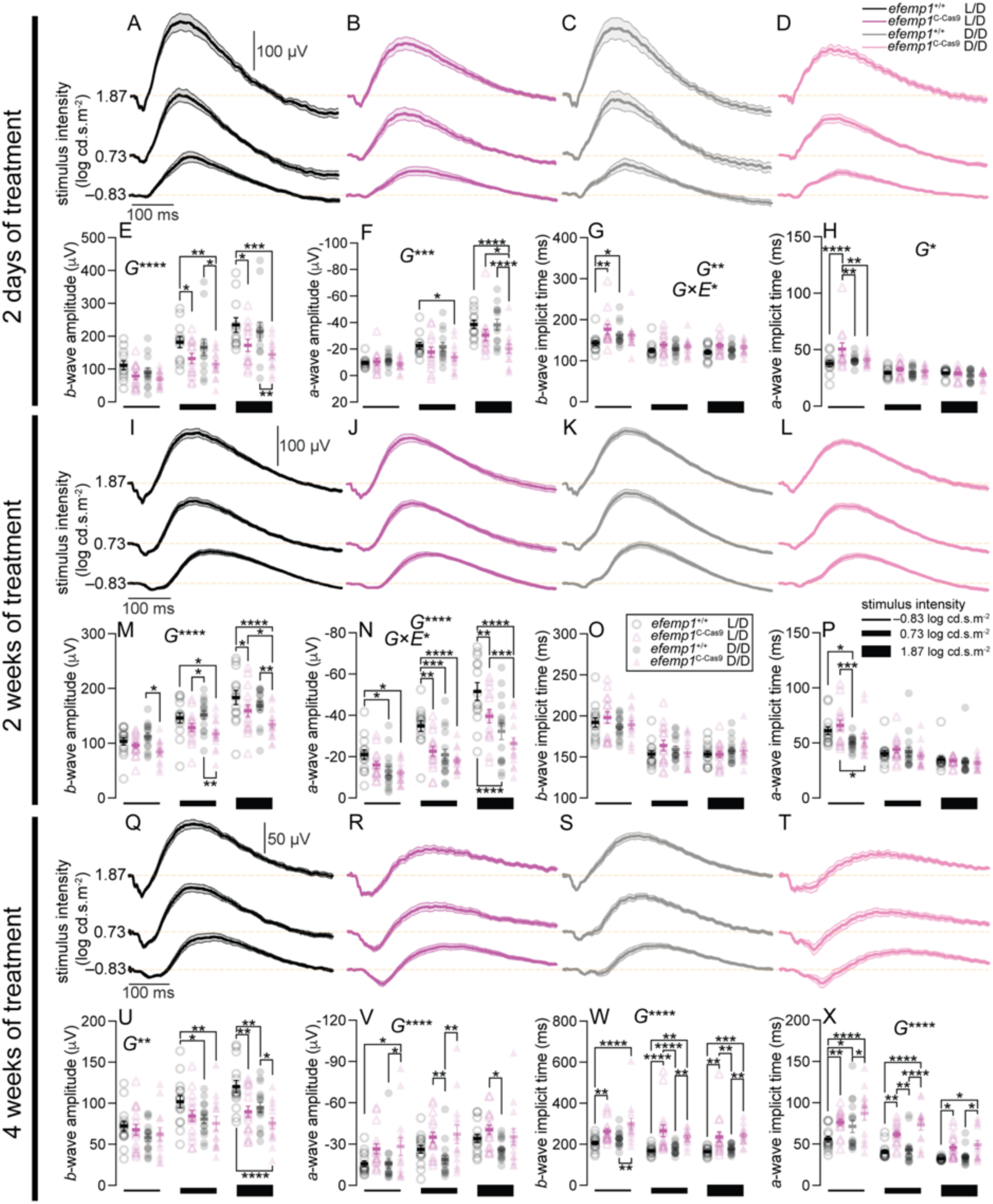
Electroretinography (ERG) of *efemp1*^+/+^ and *efemp1*^2C-Cas9^ fish after 2 days, 2 weeks or 4 weeks of light/dark-cycle (L/D) or dark (D/D) rearing. **(A–H)** Group average ERG traces for **(A)** L/D-reared *efemp1*^+/+^ (n = 12), **(B)** L/D-reared *efemp1*^2C-Cas9^ (n = 13), **(C)** D/D-reared *efemp1*^+/+^ (n = 13) and **(D)** D/D-reared *efemp1*^2C-Cas9^ eyes (n = 14). Shaded areas indicated SEM. Group **(E)** *b*-wave amplitude, **(F)** *a*-wave amplitude, **(G)** *b*-wave implicit time and **(H)** *a*-wave implicit times measured after 2 days of treatment. Vertical scale bar indicates 100 µV and horizontal scale bar represents 100 ms in **(A–D)**. **(I–P)** Group average ERG traces for **(I)** L/D-reared *efemp1*^+/+^ (n = 14), **(J)** L/D-reared *efemp1*^2C-Cas9^ (n = 15), **(K)** D/D-reared *efemp1*^+/+^ (n = 16) and **(L)** D/D-reared *efemp1*^2C-Cas9^ eyes (n = 15), as well as their **(M)** *b*-wave amplitude, **(N)** *a*-wave amplitude, **(O)** *b*-wave implicit time and **(P)** *a*-wave implicit time measured after 2 weeks of treatment. **(Q–X)** Group average ERG traces for **(Q)** L/D-reared *efemp1*^+/+^ (n = 15), **(R)** L/D-reared *efemp1*^2C-Cas9^ (n = 16), **(S)** D/D-reared *efemp1*^+/+^ (n = 16) and **(T)** D/D-reared *efemp1*^2C-Cas9^ eyes (n = 14), as well as their **(U)** *b*-wave amplitude, **(V)** *a*-wave amplitude, **(W)** *b*-wave implicit time and **(X)** *a*-wave implicit time measured after 4 weeks of treatment. Group data are shown as mean ±SEM. ERGs were measured at –0.83, 0.73 and 1.87 log cd.s.m^-2^ of stimulus intensities, represented by black blocks of different sizes below graphs; thicker blocks indicate higher intensities. *Three*-way ANOVA and Fisher’s LSD *post-hoc* tests were performed. Significant main effects of genotypes (*G*) and interactions between genotypes and rearing conditions (*G*×*E*) are indicated in the graphs. \**P* < *0.05*; \*\**P* < *0.01*; \*\*\**P* < *0.001*; \*\*\*\**P* < *0.0001*.

Functional outcomes after 2-weeks treatment (Fig 5I–L) were similar to those for 2 days of treatment. In general *b*- and *a*-wave amplitudes (*P* < *0.0001* for both; Fig 5M–N) were smaller in *efemp1*^2C-Cas9^ fish. After dark-rearing, *efemp1*^2C-Cas9^, but not *efemp1*^+/+^ fish showed further *b*-wave reduction (*P* = *0.031* at 1.87 log cd.s.m^-2^; Fig 5M). *Three*-way ANOVA returned significant interactions between genotype and dark-rearing on *a*-wave amplitude (*P* = *0.0396*), with *efemp1* disruption further attenuating dark-rearing-induced photoreceptor responses (Fig 5N). In particular, after dark-rearing, *a*-wave amplitudes were smaller at all tested intensities for *efemp1*^+/+^ fish (*P* = *0.042*, *P* = *0.0004* and *P* < *0.0001* for –0.83, 0.73 and 1.87 log cd.s.m^-2^, respectively), but only at the highest intensity for *efemp1*^2C-Cas9^ fish (*P* = *0.0009*; Fig 5N). We did not find difference in *b*-wave implicit time between groups (Fig 5O), however both *efemp1*^+/+^ and *efemp1*^2C-Cas9^ fish had slower *a*-wave implicit time after dark-rearing at –0.83 log cd.s.m^-2^ (*P* = *0.012* and *0.010*, respectively; Fig 5P). Overall, these results suggested that *efemp1* modification exacerbates functional changes seen with dark-rearing.

At the 4-week time point, under either rearing condition *efemp1*^2C-Cas9^ fish had smaller and slower *b*-waves than *efemp1*^+/+^ fish (*P* < *0.0001* and *P* = *0.002*, respectively; Fig 5Q–T, 5U and 5W,). *A*-waves in mutants were slightly larger and slower (Fig 5V and 5X, *P* < *0.0001*). The waveform indicates that slower *b*-waves expose more of the photoreceptoral response, making the *a*-wave both larger and slower.

Taken together, at 2 dpf and 2 wpf, significant statistical interactions indicate that *efemp1* modification affects the way that dark-rearing impacts retinal physiology.

### Retinal *efemp1* modification changes expression of myopia-associated genes under normal rearing and their responses to dark-rearing

In order to investigate potential molecular pathways through which retinal *efemp1* modification impacts refractive development under normal and dark-rearing, we quantified expression of a group of known myopia-associated genes using RT-qPCR after 2 days and 4 weeks of environmental manipulation. These timepoints broadly align with initiation (2 days) and consolidation (4 weeks) phases of myopia development in zebrafish (Fig 6A).

**Figure 6.**
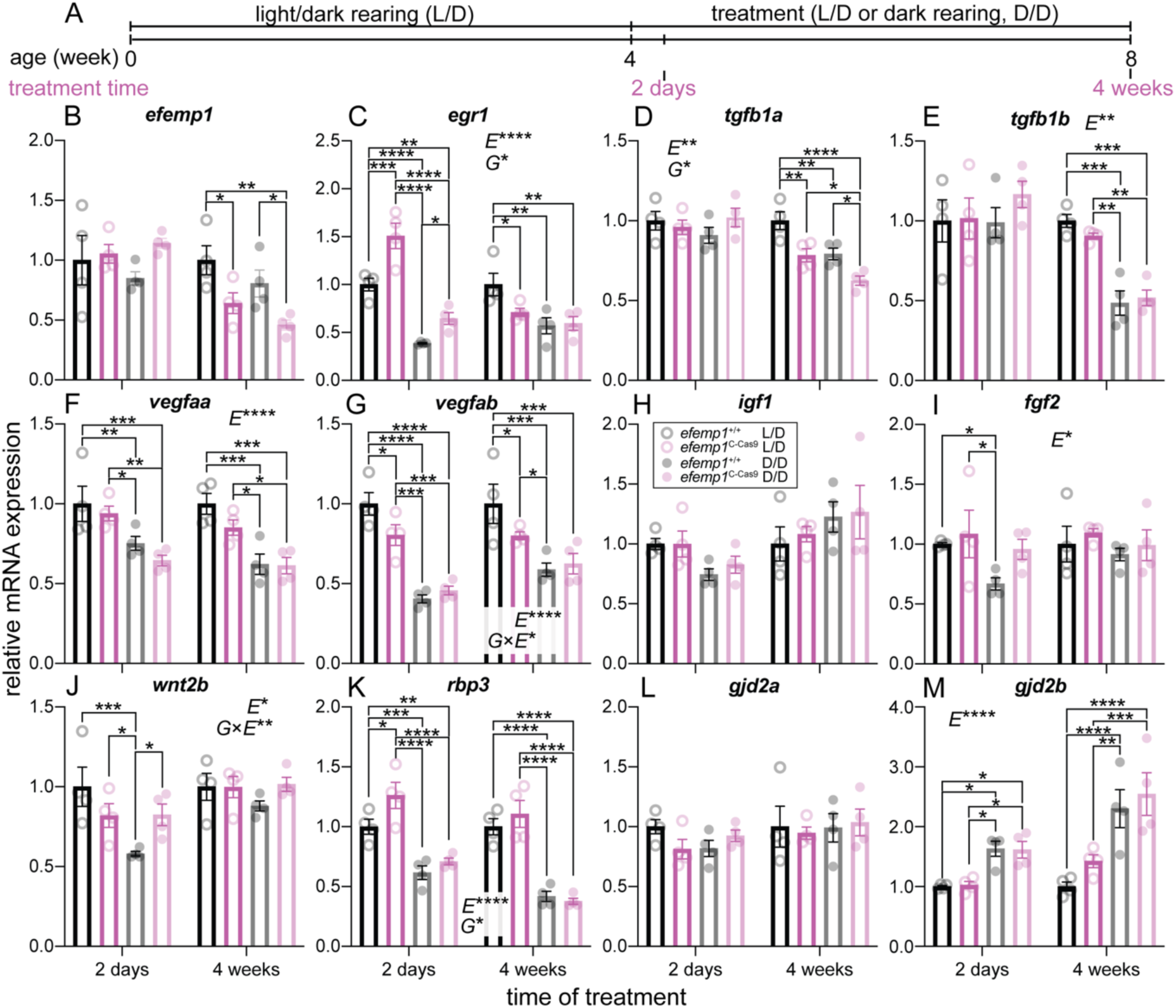
Expression of representative myopia-associated genes for *efemp1*^+/+^ and *efemp1*^2C-Cas9^ fish after 2 days or 4 weeks of light/dark-cycle (L/D) or dark (D/D) rearing. **(A)** *Efemp1*^+/+^ and *efemp1*^2C-Cas9^ fish were reared under a standard light/dark-cycle (L/D) to 4 weeks post-fertilization (wpf) and then were reared under L/D or dark tanks (D/D) to up to 8 wpf (4 weeks of treatment). Eyes for molecular analysis were sampled after 2 days or 4 weeks of treatment. Relative mRNA levels of myopia-associated genes, including **(B)** *efemp1*, **(C)** *egr1*, **(D)** *tgfb1a*, **(E)** *tgfb1b*, **(F)** *vegfaa*, **(G)** *vegfab*, **(H)** *igf1*, **(I)** *fgf2*, **(J)** *wnt2b*, **(K)** *rbp3*, **(L)** *gjd2a*, **(M)** *gjd2b* were analyzed for *efemp1*^+/+^ and *efemp1*^2C-Cas9^ fish after 2 days and 4 weeks of treatment (n = 4 samples per group), using qPCR. Group data are shown as mean ± SEM. *Three*-way ANOVA and Fisher’s LSD *post-hoc* tests were performed. Significant main effects of genotypes (*G*), rearing conditions (*E*) and their interactions (*G*×*E*) are indicated in the graphs. \**P* < *0.05*; \*\**P* < *0.01*; \*\*\**P* < *0.001*; \*\*\*\**P* < *0.0001*.

We first quantified the expression of *efemp1* and found lower *efemp1* expression in *efemp1*^2C-Cas9^ than in *efemp1*^+/+^ fish under normal and dark-rearing (*P* = *0.0248* and *0.0295*, respectively) but only at the later time point (Fig 6B).

The *early growth response 1* (*egr1*) gene is an immediate early response transcriptional factor that has been shown to be up-regulated when the eye is exposed to hyperopia-inducing or myopia-suppressing visual environments, and down-regulated when the eye is under myopia-inducing visual conditions (Ashby et al., 2014; Fischer et al., 1999). Loss of this gene promotes myopia in mice (Schippert et al., 2007). In our data, there were genotype (*P* = *0.042*) and dark-rearing effects (*P* < *0.0001*) on *egr1* expression. After 2 days of normal light rearing, *egr1* expression was up-regulated in *efemp1*^2C-Cas9^ fish compared to *efemp1*^+/+^ fish (∼ 4 wpf; *P* = *0.0002*). *Egr1* expression was then down-regulated later after 4 weeks in normal lighting (8 wpf; *P* = *0.018*). After 2 days of dark-rearing, both genotypes showed reduced *egr1* expression (*P* < *0.0001* for both). After 4 weeks of dark-rearing, only *efemp1*^+/+^ fish showed down-regulated *egr1* expression (*P* = *0.0011*) and no further change was seen in *efemp1*^2C-Cas9^ fish (Fig 6C).

*Transforming growth factor beta* genes (*tgfbs)* encode fundamental secreted TGF-βs, which has been shown to be down-regulated in form-deprived myopic tree shrew eyes to impact collagen synthesis (Jobling et al., 2004) and in particular matrix metalloproteinase 2 (MMP2) (Itoh et al., 2010). Our results showed that expression of *tgfb1a* was impacted by genotype and rearing condition (*P* = *0.0292* and *0.0070*, respectively). At the 4-week timepoint, under either normal or dark-rearing, *efemp1*^2C-Cas9^ fish showed lower *tgfb1a* expression than *efemp1*^+/+^ fish (*P* = *0.0038* and *0.0205*, respectively). *Tgfb1a* expression was reduced in both *efemp1*^+/+^ and *efemp1*^2C-Cas9^ fish after dark-rearing (*P* = *0.0051* and *0.0271*, respectively; Fig 6D). The expression of *tgfb1b* was primarily influenced by dark-rearing (*P* = *0.0047*); both *efemp1*^+/+^ and *efemp1*^2C-Cas9^ fish showed a reduction after 4 weeks of dark-rearing (*P* = *0.0003* and *0.0041*, respectively; Fig 6E).

The *vascular endothelial growth factor A* (*vegfa*) gene encodes the glycoprotein VEGFA, which contributes to vasculature development, neuronal function (Mackenzie and Ruhrberg, 2012) and extracellular matrix regulation (Kuiper et al., 2007). Decreased retinal VEGFA concentration was found in marmosets with lens-induced myopia (Zhu et al., 2021). For the two zebrafish *vegfa* paralogues, *three*-way ANOVA showed an effect of dark-rearing on expression of both *vegfaa* and *vegfab* (*P* < *0.0001* for both). Both *efemp1*^+/+^ and *efemp1*^2C-Cas9^ fish showed decreased *vegfaa* expression after 2 days (*P* = *0.0094* and *0.0026*, respectively) and 4 weeks of dark-rearing (*P* = *0.0002* and *0.012*, respectively; Fig 6F). *Vegfab* down-regulation was found in both *efemp1*^+/+^ and *efemp1*^2C-Cas9^ fish after 2 days of dark-rearing (*P* < *0.0001* and *P* = *0.0007*, respectively) but was only observed in *efemp1*^+/+^ fish at week 4 (*P* = *0.0001*). A significant interaction between genotypes and rearing conditions shown by *three*-way ANOVA analysis (*P* = *0.0125*), is indicative of a smaller dark-rearing-induced reduction in *vegfab* expression due to *efemp1* disruption. Indeed, retinal *efemp1* disruption *per se* had led to a down-regulation of *vegfab* under normal rearing (*P* = *0.039* and *0.035* at the 2-day and 4-week time points, respectively; Fig 6G).

Insulin-like growth factor 1 (IGF1), an endocrine growth hormone, is produced in the liver and then delivered to other organs, playing an important role in overall growth during development (Stratikopoulos et al., 2008). A previous study reported that *igf1* was up-regulated in form-deprived myopic guinea pigs (Ding et al., 2020). However, we did not observed difference in *igf1* expression between groups in our experiments (Fig 6H).

Fibroblast growth factor (FGF2) has functions in embryonic development, angiogenesis, wound healing, cell differentiation and neuronal function (e.g., photoreceptor transduction) (Bikfalvi et al., 1997). With form-deprivation myopia, *fgf2* expression was reported to be down-regulated in the chick sclera but up-regulated in guinea pig retinae (An et al., 2012; Seko et al., 1995). Our *three*-way ANOVA analysis showed an effect of rearing conditions on *fgf2* expression (*P* = *0.0432*). Only *efemp1*^+/+^ but not *efemp1*^2C-Cas9^ fish showed a reduction of *fgf2* expression after 2 days of dark-rearing (*P* = *0.0392*), suggesting that *efemp1* disruption may attenuate dark-rearing induced changes in *fgf2* expression (Fig 6I). Yet, our data showed no statistical significance for a gene-environment interaction.

The *Wnt family 2B* (*wnt2b*) gene activates the Wnt/β-catenin signaling pathway which is crucial for ocular development (e.g., lens, retinal pigment epithelium, vasculature and rod photoreceptors, etc.) (Blomfield et al., 2023). Mice with form-deprivation myopia were found to show up-regulated *wnt2b* expression (Ma et al., 2014). Our statistics showed a significant interaction between genotype and rearing condition (*P* = *0.0096*; Fig 6J). *Efemp1*^+/+^ fish, but not *efemp1*^2C-Cas9^ fish, showed down-regulation of *wnt2b* expression after 2 days of dark-rearing (*P* = *0.0003*), implicating reduced dark-rearing-induced changes in *wnt2b* expression with *efemp1* deficiency.

The soluble retinol binding protein 3 (RBP3; or IRBP in mammals) is released into the interphotoreceptor matrix from retinal photoreceptors, mediating retinoid trafficking for the visual cycle (Gonzalez-Fernandez and Ghosh, 2008). Lack of *Rbp3* in mice caused excessive eye growth and myopia (Wisard et al., 2011). In both *efemp1*^+/+^ and *efemp1*^2C-Cas9^ fish, *rbp3* expression was down-regulated after 2 days (*P* = *0.0007* and *P* < *0.0001*, respectively) and 4 weeks of dark-rearing (*P* < *0.0001* for both; Fig 6K). Rearing condition thus made a major effect on expression of *rbp3* (*P* < *0.0001*). Retinal *efemp1* modification also resulted in significantly higher *rbp3* expression in *efemp1*^2C-Cas9^ fish (*P* = *0.044*), particularly after 2 days under normal rearing (*P* = *0.0144*; Fig 6K).

The *gap junction delta 2* (*gjd2*) gene encodes a predominant gap junction protein, connexin36, in the central nervous system, including the retina, forming electrical synapses for rapid neuronal signaling (Bloomfield and Völgyi, 2009). Recent findings showed that depletion of the two zebrafish paralogues, *gjd2a* and *gjd2b*, caused myopia and hyperopia, respectively (Quint et al., 2021). We found no expression changes in *gjd2a* expression between groups (Fig 6L). In contrast, *three*-way ANOVA analysis showed a main effect of rearing conditions on *gjd2b* expression (*P* < *0.0001*). Both *efemp1*^+/+^ and *efemp1*^2C-Cas9^ fish showed an up-regulation after 2 days (*P* = *0.0261* and *0.0368*, respectively) and 4 weeks of dark-rearing (*P* < *0.0001* and *P* = *0.0003*, respectively; Fig 6M).

### Differences in EFEMP1, TIMP2 and MMP2 distributions between *efemp1*^+/+^ and *efemp1*^2C-Cas9^ fish were found after normal and dark-rearing

We examined whether retinal *efemp1* modification affects the distribution of myopia-associated extracellular matrix proteins in the retina after 2 days or 4 weeks of normal or dark-rearing (Fig 7A).

**Figure 7.**
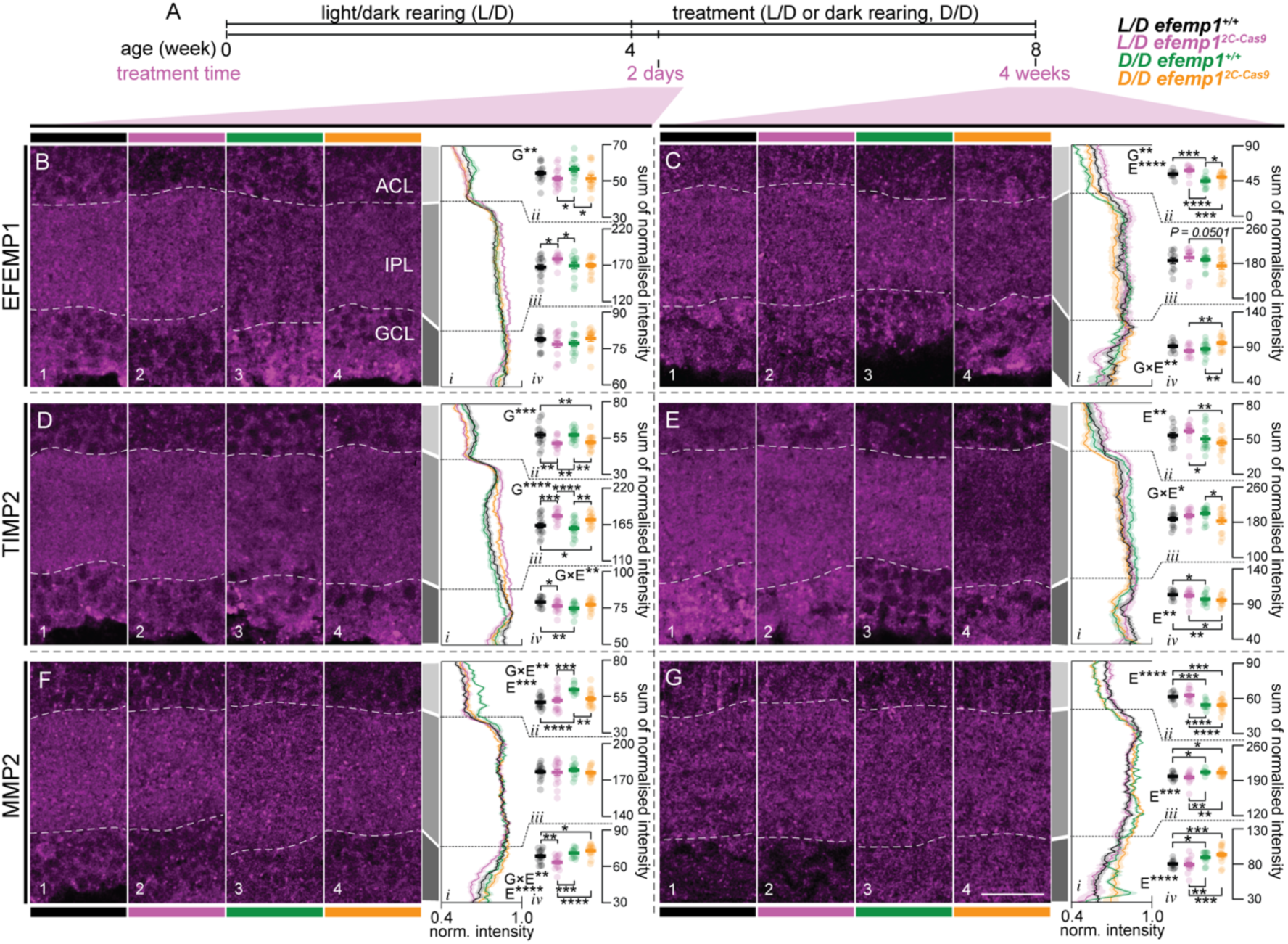
Distribution of EFEMP1, TIMP2 and MMP2 in the inner retina of *efemp1*^+/+^ and *efemp1*^2C-Cas9^ fish after 2 days or 4 weeks of normal light/dark (L/D) cycle or dark (D/D) rearing. **(A)** *Efemp1*^+/+^ and *efemp1*^2C-Cas9^ fish were reared under standard L/D condition to 4 weeks post-fertilization (wpf) and then were under L/D- or D/D-rearing to up to 8 wpf (4 weeks of treatment). Eyes for histological analysis were sampled after 2 days or 4 weeks of treatment. **(B–C)** Immunostaining for EFEMP1 to analyze its distribution in the inner retina, including amacrine cell layer (ACL), inner plexiform layer (IPL) and ganglion cell layer (GCL), for L/D-reared *efemp1*^+/+^, L/D-reared *efemp1*^2C-Cas9^, D/D-reared *efemp1*^+/+^ and D/D-reared *efemp1*^2C-Cas9^ fish after 2 days (**B**; n = 16, 16, 14 and 14 retinae, respectively) and 4 weeks (**C**; n = 12, 12, 15 and 14 retinae, respectively) of treatment. White dash lines in micrographs highlight the borders of the IPL. Average normalized expression across the inner retina is shown in (**B*_i_* and C*_i_***) and summed normalized expression (normalized intensity) for (**B*_ii_* and C*_ii_***) ACL, (**B*_iii_* and C*_iii_***) IPL and (**B*_iv_* and C*_iv_***) GCL was quantified for each eye. **(D–E)** Immunostaining for TIMP2 in the inner retina for L/D-reared *efemp1*^+/+^, L/D-reared *efemp1*^2C-Cas9^, D/D-reared *efemp1*^+/+^ and D/D-reared *efemp1*^2C-Cas9^ fish after 2 days (**D**; n = 16, 16, 16 and 15 retinae, respectively) and 4 weeks (**E**; n = 13, 14, 15 and 14 retinae, respectively) of treatment. Average normalized expression is shown in (**D*_i_* and E*_i_***) and summed normalized expression for (**D*_ii_* and E*_ii_***) ACL, (**D*_iii_* and E*_iii_***) IPL and (**D*_iv_* and E*_iv_***) GCL. **(F–G)** Immunostaining for MMP2 for L/D-reared *efemp1*^+/+^, L/D-reared *efemp1*^2C-Cas9^, D/D-reared *efemp1*^+/+^ and D/D-reared *efemp1*^2C-Cas9^ fish after 2 days (**F**; n = 16, 16, 12 and 15 retinae, respectively) and 4 weeks (**G**; n = 13, 13, 12 and 15 retinae, respectively) of treatment. Average normalized expression is shown in (**F*_i_* and G*_i_***) and summed normalized expression for (**F*_ii_* and G*_ii_***) ACL, (**F*_iii_* and G*_iii_***) IPL and (**F*_iv_* and G*_iv_***) GCL. Scale bar in **(G)** represents 20 μm and all images share the same scale. Group data are shown as mean ± SEM. *Two*-way ANOVA and Fisher’s LSD *post-hoc* tests were performed. Significant main effects of genotypes (*G*), rearing conditions (*E*) and their interactions (*G*×*E*) are indicated in the graphs. \**P* < *0.05*; \*\**P* < *0.01*; \*\*\**P* < *0.001*; \*\*\*\**P* < *0.0001*.

With the mosaic pattern of retinal *efemp1* modification (Fig 2C), EFEMP1 protein expression was still evident in the retina (Fig 7B–C). We found that at the 2-day timepoint, *efemp1*^2C-Cas9^ fish showed a change in EFEMP1 distribution with relatively lower levels in the amacrine cell layer (ACL; effect of genotypes: *P* = *0.0079*; Fig 7B_1–4_ and 7B*_i_*_–*ii*_) but higher relative expression in the inner plexiform layer (IPL) compared with *efemp1*^+/+^ fish (*P* = *0.0132*; Fig 7B_1–2_ and B*_i_*_–*iii*_). After 2 days of dark-rearing, inner retinal EFEMP1 distribution remained largely unchanged in both genotypes (Fig 7B). At the 4-week timepoint, dark-rearing reduced EFEMP1 expression in the ACL in both genotypes (effect of rearing conditions: *P* < *0.0001*; Fig 7C_1–4_ and 7C*_i_*_–*ii*_). A significant interaction was observed for the ganglion cell layer (GCL; *P* = *0.0022*; Fig 7C*_i_* and C*_iv_*), with *efemp1*^2C-Cas9^ fish (*P* = *0.0015*), but not *efemp1*^+/+^fish, exhibiting higher relative EFEMP1 expression in the GCL after dark-rearing.

Matrix metalloproteinase 2 (MMP2) and tissue inhibitor of metalloproteinase 2 (TIMP2) have long been associated with myopia (Jia et al., 2017; Jia et al., 2014; Liu and Sun, 2018) and our previous data showing that their distribution was changed after dark-rearing (Xie et al., 2023 Preprint).

Analyzing their inner retinal distribution, we found that at the 2-day timepoint, relative TIMP2 expression in *efemp1*^2C-Cas9^ fish was overall lower in the ACL but higher in the IPL, compared to control *efemp1*^+/+^ fish (effect of genotypes: *P* = *0.0001* and *P* < *0.0001*, respectively; Fig 7D_1–4_ and 7D*_i_*_–*iii*_). Dark-rearing reduced relative TIMP2 expression in the GCL for *efemp1*^+/+^ (*P* = *0.0015*) but not *efemp1*^2C-Cas9^ fish, showing a gene-environment interaction (*P* = *0.0069*; Fig 7D*_i_* and 7D*_iv_*). At the 4-week timepoint, overall dark-rearing reduced relative TIMP2 expression in the ACL and GCL (effect of rearing conditions: *P* = *0.0067* and *0.0021*, respectively; Fig 7E_1–4_, 7E*_i–ii_* and 7E*_iv_*). In the IPL, after dark-rearing, relative TIMP2 expression seemed to be reduced in *efemp1*^2C-Cas9^ fish but slightly increased in *efemp1*^+/+^ fish, exhibiting a gene-environment interaction (*P* = *0.025*; Fig 7E*_iii_*).

For MMP2, in the ACL, 2 days of dark-rearing overall resulted in higher MMP2 expression (effect of environment: *P* = *0.0008*) in *efemp1*^+/+^ fish but not in *efemp1*^2C-Cas9^ fish (gene-environment interactions: *P* = *0.0079*; Fig 7F_1–4_ and 7F*_i–ii_*). In the GCL, after 2 days of normal rearing, *efemp1*^+/+^ fish had higher relative MMP2 expression than *efemp1*^2C-Cas9^ fish (*P* = *0.0064*; Fig 7F_1–2_ and 7F*_i_*_–*iv*_). 2 days of dark-rearing overall led to increased MMP2 expression in the GCL (effect of rearing conditions: *P* < *0.0001*); this increase was greater in *efemp1*^2C-Cas9^ than in *efemp1*^+/+^ fish (effect of gene-environment interactions: *P* < *0.0075*; Fig 7F_1–4_, 7F*_i_* and 7F*_iv_*). At the 4-week timepoint, MMP2 expression pattern was affected by dark-rearing, with lower relative expression in the ACL but higher expression in the IPL and GCL (effect of environment: *P* < *0.0001*, *P* = *0.0002* and *P* < *0.0001*, respectively; Fig 7G).

Overall, inner retinal distributions of EFEMP1, TIMP2 and MMP2 can be affected by either *efemp1* disruption or dark-rearing. More importantly, retinal *efemp1* modification impacts, in a time dependent manner, how dark-rearing-affects the redistribution of inner retinal proteins.

## Discussion

In this study, we showed that retina-specific *efemp1* down-regulation leads to enlarged eyes and myopia development in zebrafish (Fig 2B, 2D–I). This resembles the progressive myopia that has been reported in human patients with loss-of-function *efemp1* mutations (Driver et al., 2020). However, mice with *efemp1* knockout showed no alternations in gross ocular structure, retinal function nor visual acuity, but did show progressive corneal maldevelopment (Daniel et al., 2020). Unfortunately, the authors did not analyze ocular refraction in their loss-of-function mouse model. In addition to aberrant eye growth, our mutants had reduced spatial visual sensitivity (Fig 1E–H, Fig S1) and altered retinal function (Figs 3 and 5). Particularly striking was the progressive slowing of the ERG in *efemp1*^2C-Cas9^ fish with older ages (Fig 3 and 5). This resembles the age-related slowing of *b*-wave implicit time reported in some, but not all, human patients with Malattia Leventinese, a retinal dystrophy caused by a gain-of-function *efemp1* mutation (c.1033C>T, p.Arg345Trp) (Gerber et al., 2003). Taken together, *efemp1* mutant zebrafish show phenotype characteristics indicative of both gain- and loss-of-function mutations.

Concurrence of gain- and loss-of-function *efemp1* mutations in our mutants could be expected; with the 2C-Cas9 system, CRISPR gene editing independently occurred in retinal neurons expressing the Cas9 nuclease, leading to different types of indels even within an individual retina. Despite these mutations being mosaic and perhaps of low efficiency (Fig 1C–D) in our model, the phenotype was robust, indicating that tight control of finely balanced *efemp1* expression is required for optimal ocular development. However, this does demonstrate the importance of using the 2C-Cas9 system with caution for studies requiring specific gain- or loss-of-function mutations. To avoid such heterogeneous tissue-specific gene editing, the Cre-LoxP system is an option: using tissue-specific driven Cre recombination to delete LoxP flanked exons of the target gene. Our data showed a down-regulation of *efemp1* in *efemp1*^2C-Cas9^ fish at 8 weeks of age (Fig 6B). In contrast, a recent study reported increased EFEMP1 in tears sampled from myopic human patients and increased EFEMP1 in myopic guinea pig choroid (Shi et al., 2023). To understand these species related differences, tissue specific comparative studies would be required.

We noticed that although 5 dpf *efemp1*^2C-Cas9^ fish overall were not myopic relative to *efemp1*^+/+^ fish (Fig 2B), they showed reduced spatial-frequency tuning function (Fig 1E–H). This phenotype, if not due to refractive error, can be a result of altered visual processing, as aberrant extracellular matrix caused by *efemp1* disruption may lead to dysfunctional synapses (Dityatev and Schachner, 2006).

For our ERG results, although slower ERG responses were a robust phenotype in *efemp1*^2C-Cas9^ fish at the older ages, at 2 wpf *a*- and *b*-wave implicit times were actually faster than control *efemp1*^+/+^ fish (Fig 3A_3_–A_4_). One possible explanation is that *efemp1*^2C-Cas9^ fish had shorter axial length than *efemp1*^+/+^fish at this age (*P* < *0.0001*, unpaired *t-*test; Fig S2) and thus effectively higher retinal illumination.

We noticed that under normal rearing, *efemp1*^2C-Cas9^ fish had larger *a*-wave amplitudes at 8 wpf (or 4-week timepoint in the dark-rearing experiment) than *efemp1*^+/+^ fish, which we attributed to slower *b*-wave onset exposing more of the *a*-wave (Fig 3D_2_ and Fig 5V). *B-wave* effects in 8 wpf *efemp1*^2C-Cas9^ fish were less consistent, with slightly higher *b*-wave amplitudes in one experiment (Fig 3D_1_) but smaller *b*-waves in another (Fig 5U). We speculated that this might arise from variability in gene editing (as discussed above) between retinae (and batches), leading to different severities and phenotypes. Nonetheless, it is clear that *efemp1* function causes retinal dysfunction in zebrafish.

Under dark-rearing, retinal dysfunction in *efemp1*^2C-Cas9^ was exacerbated (Fig 5), but this did not lead to worsening myopia (Fig 4). One speculation is that without visual input (dark-rearing) in the presence of *efemp1* disruption the zebrafish eye shuts down light-dependent signals for ocular growth and switches to intrinsic mechanisms for ocular growth regulation, thus leading to emmetropic vision. This idea requires investigation.

Our survival analysis suggested that *efemp1*^2C-Cas9^ mutants at 4 wpf did not show a myopic shift. This could be due to a number of factors. Firstly, because those fish with a higher level of visual impairment and perhaps early-onset myopia are those likely to have been lost by 4 wpf. Thus, those fish with lower mutation rates and better vision were more likely survived to older ages allowing more time (>4 weeks) for the development of myopia (Fig 2C). Secondly, changes in *egr1* expression shown in our data (Fig 6C) may also be a separate contributor to the observed absent myopic shift in 4 wpf *efemp1*^2C-Cas9^ fish. Previous studies suggest that *egr1* may have a role in signaling bi-directional ocular growth (Ashby et al., 2014; Fischer et al., 1999). Up-regulated *egr1* expression in 2-day normal-light-reared (∼4 wpf) *efemp1*^2C-Cas9^ fish showed evidence myopia suppression, whereas down-regulation of *egr1* in 4-week normal-light-reared (equivalent to 8 wpf) *efemp1*^2C-Cas9^ fish showed a myopic shift (Fig 6C). The direction of these expression changes qualitatively matched ocular refraction of *efemp1*^2C-Cas9^ fish at the corresponding ages (Fig 2B).

In our gene expression data, down-regulation of *tgfb1a* and *vegfab* was associated with myopia development in *efemp1*^2C-Cas9^ fish under normal lighting (Fig 6E and 6G) (Jobling et al., 2004; Zhu et al., 2021). Dark-rearing-induced down-regulation of *egr1*, *tgfb1s*, *vegfaa*, *fgf2* and *rbp3*, and up-regulation of *gjd2b* were not impacted by *efemp1* modification (Fig 6C, 6D–F, 6I, 6K and 6M), with two exceptions being *vegfab* and *wnt2b* (Fig 6G and 6J). Despite the similarities in dark-rearing-induced gene expression, *efemp1*^+/+^ fish became myopic after 4 weeks of dark-rearing, *efemp1*^2C-Cas9^ fish remained emmetropic (Fig 4). It is possible that *efemp1* functional integrity is required to manifest dark-rearing-induced phenotypes with changes in these genes. We speculate that without normal *efemp1* expression under dark-rearing, intrinsic mechanisms may take over and regulate ocular growth. Future investigations are required to verify this hypothesis.

The genes *wnt2b* (Fig 6J) and *vegfab* (Fig 6 G), were down-regulated only in *efemp1*^+/+^ but not *efemp1*^2C-Cas9^ fish after 2 days and 4 weeks of dark-rearing, respectively, suggesting that dark-rearing requires normal *efemp1* expression to induce and maintain their expression changes, respectively. EFEMP1 has been previously shown to suppress the epithelial-mesenchymal transition in endometrial carcinoma through Wnt/β-catenin signaling, for which WNT2B is an activator (Blomfield et al., 2023; Yang et al., 2016). Further investigation might target interactions between EFEMP1, WNT2B and EMT in myopia development. Importantly, the impact of *efemp1* modification on dark-rearing-induced *vegfab* expression further implies a light-dependent signaling between *efemp1* and *vegfab* that is crucial for ocular growth.

There was up-regulation of *rbp3* expression in normal-light-reared *efemp1*^2C-Cas9^ fish at the 2-day but not at the 4-week timepoint (Fig 6K). It is likely that *efemp1* deficiency altered the interphotoreceptor matrix and thus impairing the visual cycle and *11-cis* retinal supply (Gonzalez-Fernandez and Ghosh, 2008; Yue, 2014). Up-regulating *rbp3* may be a compensatory response in young zebrafish. Myopia development has been reported in *gjd2b* knockout zebrafish (Quint et al., 2021), however our data showed robust *gjd2* up-regulation in both phenotypes under myopia-inducing dark-rearing (Fig 6M). Whether this up-regulation reflect feedback remains to be clarified.

In our histological analysis, distribution changes of EFEMP1 in the inner retina under different rearing conditions indicate retinal *efemp1* modification at the post-transcriptional level (Fig 7B–C). In addition, our data indicated that *efemp1* modification affects the way that inner retinal MMP2 (particularly after 2 days) and TIMP2 (particularly after 4 weeks) were impacted in response to dark-rearing (Fig 7D–G). There have been previous reported that changes in expression of EFEMP1 in cultured endothelial cells led to decreased MMP2 but unchanged TIMP2 expression (Albig et al., 2006), highlighting a context-dependent regulation of EFEMP1. In future studies, it will be worth analyzing other MMPs (e.g., MMP3, etc.) and TIMPs (e.g., TIMP1 and TIMP3, etc.) that have been associated with EFEMP1 (Livingstone et al., 2020) and eye elongation (Jia et al., 2017; Jia et al., 2014).

In summary, the modification of retinal *efemp1* in zebrafish resulted in reduced spatial visual sensitivity, axial myopia, eye enlargement, altered retinal function, myopia-associated gene expression changes (i.e., *egr1*, *efemp1*, *tgfb1a*, *vegfab*, *rbp3*), and redistribution of inner retinal EFEMP1, TIMP2 and MMP2 proteins, under normal rearing. Under dark-rearing, *efemp1*^2C-Cas9^ fish did not develop myopia, even though dark-rearing seemed to worsen retinal function. We found *efemp1*-dependent responses of *vegfab* and *wnt2b* expression to dark-rearing, highlighting signaling between *efemp1* and these genes as likely mechanisms involved in visually regulated ocular development. Dark-rearing-induced redistribution of EFEMP1, TIMP2 and MMP2 were also influenced by *efemp1* disruption. This study provides robust evidence that *efemp1* regulates ocular development and visual function in a light-dependent manner, with several potential myopia-associated molecular pathways implicated for further investigation. We illustrated a valuable zebrafish analysis platform combining environmental and genetic manipulations, as well as multifaced analyses for high-throughput investigation of the growing number of myopia-risk genes to gain insight into their interactions with visual environment in myopia development.

## Materials and Methods

### Animal husbandry

All procedures were performed according to the provisions of the Australian National Health and Medical Research Council code of practice for the care and use of animals and were approved by the Faculty of Science Animal Ethics Committee at the University of Melbourne (Project No. 10399).

For standard husbandry, zebrafish (Danio rerio; transgenic fish, see detailed description below) were maintained and bred in the Fish Facility at the University of Melbourne according to local animal guidelines. Embryos and larvae (prior to sex determination) were grown in Petri dishes in an incubator at 28.5°C up to 5 days post-fertilization (dpf), then introduced to tanks under normal 14h/10h light/dark (L/D) cycles and raised in flow-through systems at 28°C.

For dark (D/D) rearing, zebrafish tanks were wrapped using black cloth tapes (Model number 66623336603; Bear brand, Saint-Gobain Abrasives, Somerton, VIC, Australia) to block light. Zebrafish were induced to blackout tanks at 4 wpf and reared in darkness for 2 days, 2 or 4 weeks. Age-matched fish reared under standard lighting were used as environmental control. Zebrafish health condition and survival were monitored daily using dim red light (LED; 17.4 cd.m^-2^, λ_max_ 600 nm).

### Generation of mutants

In this study, we used a novel somatic CRISPR gene editing system, namely 2C-Cas9 (Di Donato et al., 2016), to generate retina-specific *efemp1* mutation in zebrafish. The p*UAS*:*Cas9T2ACre*;*U6*:*sgRNA1*;*U6*:*sgRNA2* plasmid (Addgene #74010) was digested using BsaI (R0535S; NEB, Ipswich, MA, USA) and BsmBI-v2 enzymes (R0739S; NEB) targeting the restriction sites behind the U6 promoters to remove the pre-set sequences that are to be replaced by DNA sequences of small guide RNA (sgRNA). Digested vectors were purified using a QIAquick Gel Extraction kit (28704; Qiagene, Hilden, Germany). Double-strand DNAs of the two sgRNAs (oligos shown in Table S2), *efemp1sgRNA1* and *efemp1sgRNA2*, were annealed from single-strand oligos and inserted into the digested vector using T4 DNA ligase (M0467S; NEB). Ligated plasmids were transformed into α-select chemically competent cells (BIO-85026; Bioline, London, UK). Positively transformed cells were selected using antibiotic Ampicillin LB agar plates and colonies were picked to grow in Ampicillin LB medium at 37°C for overnight. Plasmids were extracted and purified using a QIAprep Spin Miniprep kit (27106; Qiagene). Successful insertions of DNAs into the plasmid were determined by sequence-specific PCR, gel electrophoresis and Sanger sequence (Fig S3B). Concentrations of extracted plasmids were quantified using a Nanodrop 1000 (Thermo Fisher Scientific; Waltham, MA, USA). *Efemp1* sgRNAs targeting exons were designed based on the *efemp1* genome DNA sequence (gDNA; ENSDARG00000059121) using the *CHOPCHOP* online tool (Labun et al., 2019) and pre-tested *in vitro* using a GenCrispr sgRNA Screening kit (Fig S3A;L00689; GenScript, NJ, USA) before insertion into vectors. The two sgRNAs target exons 3 and 5 of *efemp1* gDNA. Tol2 transposase mRNA was transcribed from a p*tol2* plasmid (kind gift from Dr. Mirana Ramialison, Murdoch Children’s Research Institute) using a mMESSAGE mMACHINE SP6 Transcription Kit (AM1340; Thermo Fisher Scientific).

To generate the retina-specific *efemp1* modification in zebrafish, we co-injected p*UAS*:*Cas9T2ACre*;*U6*:*efemp1sgRNA1*;*U6*:*efemp1sgRNA2* at a concentration of 35 ng/μl and *tol2* mRNA at 50 ng/μl into embryos of fish Tg(*bact2*-*loxP*-*mCherry*-*loxP*-*eGFP*) × Tg(*rx2*:*Gal4*). Expression of the injected plasmid were identified based on green fluorescence (eGFP) in the zebrafish eyes observed under a fluorescence microscope at 3 dpf. F0 fish with green fluorescent eyes were mated with transgenic fish Tg(*bact2*-*loxP*-*mCherry*-*loxP*-*eGFP*) × Tg(*rx2*:*Gal4*) to screen for the founders that had the 2C-Cas9 transgene integrated into the germ line. In this study, we used transgenic fish Tg(*bact2*-*loxP*-*mCherry*-*loxP*-*eGFP*) × Tg(*rx2*:*Gal4*), namely *efemp1*^+/+^ fish as the control of the mutant fish Tg(*bact2*-*loxP*-*mCherry*-*loxP*-*eGFP*) × Tg(*rx2*:*Gal4*) × Tg(*UAS*:*Cas9T2ACre*;*U6:efemp1sgRNA1*;*U6:efemp1sgRNA2*), named *efemp1*^2C-Cas9^ fish. These fish also have a transgene Tg(*UAS*:*nfsb*-*mCherry*) that was driven by the retina-specific Tg(*rx2*:*Gal4*), leading to much brighter red fluorescence signals in the zebrafish eye than in the body, where *mCherry* expression was triggered by Tg(*bact2*-*loxP*-*mCherry*-*loxP*-*eGFP*). These allowed to sort Tg(*rx2*:*Gal4*) positive fish for *efemp1*^+/+^ fish. *Efemp1*^2C-Cas9^ fish were sorted by eye-specific green fluorescence, which indicated presence of Tg(*rx2*:*Gal4*), without the need to the Tg(*UAS*:*nfsb*-*mCherry*) reporter. However, to ensure that the only genetic difference between *efemp1*^+/+^ and *efemp1*^2C-Cas9^ fish was from the 2C-Cas9 construct Tg(*UAS*:*Cas9T2ACre*;*U6:efemp1sgRNA1*;*U6:efemp1sgRNA2*), we also used fish with bright red eyes indicating existence of Tg(*UAS*:*nfsb*-*mCherry*) for mutant groups. As the Tg(*UAS*:*nfsb*-*mCherry*) was not directly relevant to the 2C-Cas9 gene editing system and our phenotypic assessments, for a simplified demonstration, we did not highlight this transgene construct in this report.

For genotyping, we used a headloop PCR protocol (Kroll et al., 2021). Standard PCR Primers were designed using Primer Premier 5.0 (Singh et al., 1998) and headloop PCR primers were designed using a published Python script (Kroll et al., 2021) (see Table S3 for oligo information). GDNA was isolated from 7 dpf *efemp1*^+/+^ or *efemp1*^2C-Cas9^ zebrafish eyes using a Monarch Genomic DNA Purification Kit (T3010S; NEB) and quantified using a Nanodrop 1000 (Thermo Fisher Scientific). In a PCR reaction, there were 30 ng gDNA of either genotype, 0.25 μl 10 mM forward primer, 0.25 μl 10 mM reverse primer, 5 μl Hot Start High-Fidelity 2× Master Mix (M0494S; NEB) and nuclease-free water to fill up to 10 μl. The PCR program was comprised of an initial denaturation step of 98°C for 30 seconds, 32 cycles of 98°C for 15 seconds, 60°C for 15 seconds, and 72°C for 15 seconds, followed by a final extension step of 72°C for 2 minutes. Electrophoresis of the PCR products was performed using 2% Tris-acetate-EDTA (TAE) agarose gel.

### Optomotor response (OMR)

The OMR apparatus was adapted from that previously described (Xie et al., 2019a; Xie et al., 2021; Xie et al., 2019b). A Power Mac G5 computer (Apple Computer, Inc., Cupertino, CA, USA) ran MATLAB 2016b (MathWorks, Natick, MA, USA) with Psychtoolbox extensions (Kleiner et al., 2007). Stimuli were processed on an ATI Radeon HD 5770 graphics card, with the output sent to a BITS++ video processor (Cambridge Research Systems, Rochester, UK) for increased contrast resolution for stimuli displayed on a M992 flat-screen cathode ray tube (CRT) monitor (Dell Computer Corporation, Round Rock, TX, USA) with its screen facing upwards. During experiments, a zebrafish was placed in a custom annulus swimming chamber (the inner and outer walls were 29.5 and 46.5 mm from the center, respectively) with a transparent base positioned 46 mm above the screen. A C922 Pro Stream webcam (Logitech Company, Lausanne, Switzerland; 1080p at 30 Hz), placed 130 mm above the swimming chamber base, controlled using MATLAB recorded videos of each trial for *post-hoc* analysis of zebrafish angular movement.

During a trial, a test stimulus was displayed on the CRT screen below the swimming chamber, rotating at 0.5 rad/s for 30s. Test stimuli were windmill sinusoidal gratings with central spatial frequencies of 0.0078, 0.0155, 0.0310, 0.0620, 0.1240 and 0.2480 cycle per degree (c/°). A blank grey screen was presented between test stimuli. Test stimuli were generated using the green and blue channels of the CRT monitor with constant red luminance across the screen. A long pass filter (cut-off wavelength 600nm) was fixed to the front of a camera to filter out blue and green light, allowing only red light to pass through, thus eliminating the moving stimulus and allowing better visualization of the fish for analysis.

OMR was measured for 5 dpf and 2 wpf *efemp1*^+/+^ and *efemp1*^2C-Cas9^ zebrafish, after acclimating fish to the swimming chamber for 5 minutes. All experiments were conducted between 9:00 AM and 7:00 PM. After experiments, fish were humanely killed using 1000 ppm AQUIS (#106036, Primo Aquaculture).

The position of zebrafish during each trail was tracked using a published python package *Stytra* (Štih et al., 2019). Using custom MATLAB algorithms, we calculated the overall angular movement of zebrafish in the direction of the rotating grating for each trial as the optomotor index (OMI). For each fish, OMI at a spatial frequency was averaged from results of 4 trials (2 repeats at either clockwise or counter-clockwise direction). 5 dpf OMI data were normalized to the averaged OMI of *efemp1*^+/+^ fish at 0.0155 c/° (i.e., the condition for which 5 dpf zebrafish showed the greatest response) and 2 wpf OMI data were normalized to averaged OMI of *efemp1*^+/+^ fish at 0.0620 c/° (i.e., the condition for which 2 wpf zebrafish showed the greatest response). The spatial-frequency tuning functions were fit using a log-Gaussian using a least-squares criterion to return amplitude (i.e., height of the peak), peak spatial frequency (i.e., spatial frequency at which amplitude peaked), and bandwidth (i.e., standard deviation).

To test whether spatial-frequency tuning functions differed between groups, an omnibus *F*-test was used to compare the goodness of fit (*r*^2^) of a full model, in which parameter estimates of each group could vary independently, with that of a restricted model, in which parameters were constrained to be the same across groups. To determine whether specific parameter estimates differed between groups, a nested *F*-test was used to compare a full model with a restricted model in which one parameter was constrained to be the same across groups (Lu and Dosher, 2013). A criterion of α = 0.05 was used to determine significance.

### Optical coherence tomography (OCT)

OCT was performed immediately after zebrafish were humanely killed using 1000 ppm AQUI-S (#106036, Primo Aquaculture) in E3 medium. Zebrafish were positioned on a square (2 × 2 cm) of paper towel and transferred onto a moistened PVA sponge fitted snugly in a 35-mm Petri dish. All OCT images were recorded using a spectral domain OCT/OCTA system (Spectralis OCT2; Heidelberg Engineering, Heidelberg, Germany) with the aid of a Digital High Mag lens (78D, Volk Optical Inc., Mentor, Ohio, USA) attached. The Petri dish was then affixed vertically onto a platform, allowing the zebrafish eye to be aligned with the lens. Surface tension between the moistened sponge/paper towel and the fish kept it in place. Images were acquired in the dorsal-ventral direction with a volume scan pattern of 15° × 5° (2.7 × 0.9 mm). Each volume consisted of 128 B-scans (5 repeats) each consisting of 512 A-scans. When imaging, the OCT was positioned such that the iris was horizontal, and a bright reflection can be seen at the apex of the cornea. This ensured that the image captured was aligned with the central axis of the eye. As myopia can independently develop in an eye without signals from the brain (Troilo et al., 1987), and thus the other eye, both eyes were imaged for each zebrafish in our study.

Only for 5-dpf fish, eyes were imaged using a Bioptigen Envisu R2200 spectral domain OCT system with an 18-mm telecentric lens (Bioptigen Inc., Durham, NC), due to temporary unavailability of the Spectralis OCT system. The position of the sponge platform with a zebrafish (as described above) was carefully adjusted to ensure that the central apex was aligned with the objective lens. A volume scan of 2 × 2 mm was acquired with a depth of 1.7 mm (1000 A-scans per B-scans).

Image quantification was performed using a custom MATLAB script. Axial length is quantified as the distance from the corneal apex to the retinal pigment epithelium (RPE). Lens radius is half of the distance from the anterior to posterior lens surface. The retinal radius is the distance from the center of the lens to the RPE, which indeed can be calculated as the difference between axial length and lens radius (Fig 2A). With these parameters, relative axial ocular refraction is given by the ratio of retinal radius to lens radius (*Matthiessen’s Ratio*), which is commonly used for analyzing ocular refraction of aquatic species (Collery et al., 2014; Shand et al., 1999; Turnbull et al., 2015). *Two-* or *three*-way ANOVA with Fisher’s LSD tests were performed in Prism 9 (GraphPad, San Diego, CA, USA) and a criterion of α = 0.05 was used to determine statistical significance.

### Survival comparison between *efemp1*^2C-Cas9^ fish with different eGFP levels

*Efemp1*^2C-Cas9^ fish were sorted under a fluorescence microscope and divided to two groups based on the intensity of the retinal eGFP fluorescence; fish with weak eGFP signals were classified to eGFP+ group and those with strong eGFP fluorescence were categorized to the eGFP+++ group. *Post-hoc* retinal histology indicated that intensity of eGFP fluorescence is corresponding to eGFP positive cell number; fish with higher eGFP fluorescence level had more eGFP positive cells. Both groups were reared under standard husbandry and their survival was recorded at 2, 4, 6 and 8 weeks of age. There were 3 tanks for both groups (37–60 fish each tank at 5 dpf). *Two*-way ANOVA and Fisher’s LSD *post-hoc* analyses were performed in Prism 9. A criterion of α = 0.05 was used to determine significance.

### Electroretinography (ERG)

The procedure for scotopic ERG was adapted from our published method (Xie et al., 2019c). Fish were dark adapted (>8 hours, overnight) prior to experiments. All procedures were conducted under dim red illumination (17.4 cd·m^−2^; λmax = 600 nm). For recording, a fish was humanely killed by immersing in 0.1% tricaine MS-222 (E10521; Sigma-Aldrich, St. Louis, MO, USA) in E3 medium. When gills had stopped moving and fish were unresponsive response to a gentle touch, the head was removed and immediately rinsed in E3 medium. The eyes were quickly dissected using fine-tipped tweezers under a dissecting microscope. A small incision was made around the central cornea for each eye to allow a small amount of aqueous humor to flow out. This helps keep the recording electrode to stay moist, increasing conductivity and minimizing noise. The dissected eyes were moistened with multiple drops of E3 medium using a 3-mm Pasteur pipette. The eyes were then transferred onto a square of paper towel (2 × 2 cm) on a moist polyvinyl alcohol (PVA) sponge platform in a Faraday cage. Both eyes were measured at the same time. For recording from eyes at or older than 4 wpf, recording electrodes were 0.3-mm chloride-electroplated silvers (99%). For younger eyes, a moistened cone-shape sponge (tip diameter ∼40 μm) was attached to the silver. Under a microscope, the recoding electrodes were positioned to gently touch the central corneal apex, and the reference electrodes were inserted into the sponge platform. After electrode placement, the Ganzfeld bowl was moved to cover the platform, and the eyes were allowed to dark adapt for 3 minutes. ERG responses for each fish were recorded with flash stimuli of –2.11, –0.81, 0.72, 1.89 and 2.48 to analyze the impact of *efemp1* modification on retinal function during normal development, and at −0.83, 0.73 and 1.87 log cd·s·m^−2^ for examining how *efemp1* modification impact fish’s retinal physiological changes in response to myopia-inducing dark-rearing. For stimulus intensities at or lower than 0.73 log cd·s·m^−2^, three repeats were measured with an inter-flash interval of 15 seconds and there was a 30-second interval between intensities. For stimulus intensities at or higher than 1.87 log cd·s·m^−2^, a single response was measured with a 60-second re-adaption to dark before the flash.

All experiments were performed between 9:00 AM and 7:00 PM at room temperature. Amplitudes of the *a*- and *b*-waves were measured from baseline to the negative *a*-wave trough and from the negative *a*-wave trough to the *b*-wave peak, respectively. Implicit times of the *a*- and *b*-waves were measured from stimulus onset to the *a*-wave trough and the *b*-wave peak, respectively. *Two-* or *three*-way ANOVA, and Fisher’s LSD *post-hoc* tests were performed in Prism 9. A criterion of α = 0.05 was used to determine significance.

### Gene expression analysis

Zebrafish were humanely killed by 1000 ppm AQUIS (#106036, Primo Aquaculture) and stored in RNALater (AM7020, Thermo Fisher, Waltham, MA, USA) at –20°C if not immediately used for RNA isolation. For RNA isolation, eyes were dissected from zebrafish and lenses were removed. For each sample, the total RNA of 4 eyes from two fish was isolated using TRIzol (or TRI-reagent; AM9738, Thermo Fisher scientific) (Rio et al., 2010) and quantified using a Nanodrop 1000 (Thermo Fisher Scientific). 750 ng (2 days of treatment) or 1 μg (4 weeks of treatment) of the total RNA was reverse-transcribed into cDNA using a Tetro cDNA synthesis kit (BIO-65042, Bioline). Quantitative PCR (RT-qPCR) assays were performed on a CFX96 real-time PCR system (Bio-Rad, Hercules, CA, USA) using SsoAdvanced Universal SYBR Green Supermix (12 μL final mix per reaction; 1725272, Bio-Rad). In this study, mRNA levels of *egr1*, *efemp1*, *tgfb1a*, *tgfb1b*, *vegfaa*, *vegfab*, *igf1*, *fgf2*, *wnt1b*, *rbp3*, *gjd2a* and *gjd2b* were quantified. Primers of the tested genes were as listed in our previous study (Xie et al., 2023 Preprint). A housekeeping gene *elongation factor 1-alpha* (*ef1a*) served as an internal control (McCurley and Callard, 2008). For each gene, there were 4 samples per group and 3 replicates per sample. Calculation of expression levels was performed using the 2^-ΔΔCT^ method (Livak and Schmittgen, 2001) and the transcript levels were normalized to the *ef1a* transcript levels in L/D-reared *efemp1*^+/+^ eyes at each time point. For statistics, *two*-way ANOVA and Fisher’s LSD *post-hoc* tests were performed in Prism 9. A criterion of α = 0.05 was used to determine significance.

### Histology

For Immunohistochemistry, whole zebrafish (if at 2 weeks of age) or dissected zebrafish eyes (if older than 2 weeks of age; lenses excluded) were fixed in 4% paraformaldehyde (PFA) in PBS for 48 hours at 4°C. They were cryoprotected in 30% sucrose in PBS, embedded in Tissue-Tek OCT compound (Sakura Finetek, Torrance, CA, US) and cryosectioned (20 μm of thickness; CM 1860 Cryostat; Leica, Wetzlar, Germany). Antibody staining was carried out at room temperature using standard protocols. Slides were blocked in 5% fetal bovine serum (FBS) for 30 minutes and incubated overnight in rabbit anti-EFEMP1 (ARP41450_P050, 1:600; Sapphire Bioscience, NSW, Australia), sheep anti-TIMP2 (1:1000; kind gift from the Itoh Lab at the University of Oxford, UK) or rabbit anti-MMP2 (AS-55111, 1:250; AnaSpec, Fremont, CA, USA) primary antibodies diluted in FBS. Noting that the anti-EFEMP1 antibody does not bind to EFEMP1 regions corresponding to sgRNAs target sites on zebrafish *efemp1* DNA, thus mutant proteins (if any) may still be labeled by the antibody. The anti-TIMP2 antibody was developed based on human TIMP2. In a previous study this antibody was showed to label for zebrafish TIMP2a (Zhang et al., 2003). As similarity of zebrafish TIMP2b to human TIMP2 is much lower than that of zebrafish TIMP2a (60.55% vs. 71.23%), labelling of zebrafish TIM2b is less likely. Yet, we are not able to completely rule out this possibility due to lack of information of the exact immunogen. Slides were subsequently incubated for 2 hours in secondary antibodies (all 1:500; Thermo Fisher Scientific) diluted in FBS. The secondary antibodies used were Chicken anti-Rabbit Alexa Fluor 647 (A21443) or Donkey anti-Sheep Alexa Fluor 647 (A21448). Nuclei were counterstained with 4′, 6-diamidino-2-phenylindole (DAPI; D9542-10MG, 1:10000; Sigma-Aldrich) in PBS for 20 minutes, and sections were mounted in Mowiol (81381-250G; Sigma-Aldrich). Stained sections were imaged at a distance 100 µm from the center of the optic nerve, using a Nikon A1R confocal microscope (Nikon, Minato City, Tokyo, Japan) with a 40× air objective lens (NA 0.95). For all confocal imaging, the deconvolution function of the NIS elements software (AR4.6, Nikon) was applied to minimize background noise. Image brightness and contrast were adjusted using Photoshop (Adobe, San Jose, CA, USA) or FIJI (Schindelin et al., 2012).

To analyze the relative distribution of EFEMP1, TIMP2 and MMP2, images were pre-processed in FIJI. In each image, an inner retinal region was selected by drawing a line that followed the contour of the inner plexiform layer (IPL) with a line thickness of 1000 pixels, which includes the inner nuclear layer (INL; including the amacrine cell layer, ACL), IPL and ganglion cell layer (GCL). This line allowed the retinal cross section to be straightened using the ‘Straighten’ function. The duplicated images returned were rotated, positioning the INL on the left and GCL on the right. The widths of the straightened images were then resized using Adobe Photoshop to normalize the IPL thickness to the averaged IPL thickness of the control group (L/D-reared *efemp1*^+/+^ fish) relevant to that timepoint. Regions of interest (ROI) boxes measuring 150-pixel height and widths of 390 (2-day L/D- or DD- reared *efemp1*^+/+^ and *efemp1*^2C-Cas9^ fish) or 455 pixels (4-week L/D- or DD-reared *efemp1*^+/+^ and *efemp1*^2C-Cas9^ fish) were used to return expression levels using the ‘*Plot Profile*’ function in FIJI. For each image, brightness profiles across the ACL, IPL and GCL were normalized to the highest value. To examine relative change in the three layers, normalized intensity was summed throughout the ACL (pixel 1–90), IPL (2-day: pixel 91–300; 4-week: pixel 91–330) and GCL (2-day: pixel 301–390; 4-week: pixel 331–455) for each image. One section per retina was used and both eyes from a zebrafish were taken for analysis in this study. *Two*-way ANOVA and Fisher’s LSD *post*-*hoc* tests were performed in Prism 9 and a criterion of α = 0.05 was used to determine statistical significance.

## Data availability

All data have been made available on Open Science Frame (https://osf.io/rdmvy/).

## Acknowledgments

We thank Yoshi Itoh for his kind gift of a sheet anti-TIMP2 antibody and Dr. Mirana Ramialison for providing the p*Tol2* plasmid. We thank the Biological Optical Microscopy Platform (BOMP) at the University of Melbourne for providing instrument for confocal imaging. We acknowledge the staff of the Danio rerio University of Melbourne, and Walter and Eliza Hall of Medical Institute fish facilities for animal husbandry and support.

JX was supported by the Melbourne Research Scholarship for his PhD study the Dr Albert Shimmins Postgraduate Writing-Up Award for manuscript preparation. Funding was provided through the University of Melbourne research grant support scheme and an international collaborative fund from Research Centre for SHARP vision (RCSV), The Hong Kong Polytechnic University.

## Author Contributions

Conceptualization, JX, BVB, PTG, PJ; Methodology, JX, BVB, PTG, PJ; Data collection, JX; Writing – Original Draft, JX; Writing – Review & Editing, JX, BVB, PJ; Data analysis and visualization, JX; Supervision, BVB, PTG, PJ; Project Administration, PJ; Funding Acquisition, PJ.

