## Supplementary material for "EFEMP1 contributes to light-dependent ocular growth in zebrafish"

This document includes:

Figure S1–S3

Table S1–S3

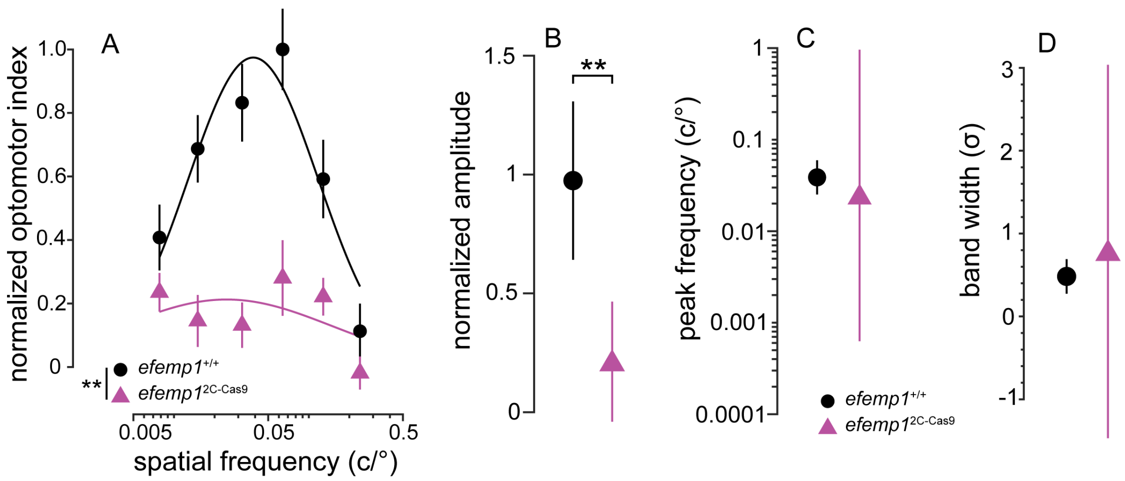


Figure S1. **Optomotor responses of *efemp1*$^{+/+}$ and *efemp1*$^{2C\text{-}Cas9}$ fish at 2 weeks post-fertilization (wpf). (A)** Spatial-frequency tuning functions are three-parameter log-Gaussian functions fit to the data by minimizing the least-square error. There were 18 *efemp1*$^{+/+}$ and 17 *efemp1*$^{2C\text{-}Cas9}$ fish used for OMR. Group data are shown as mean ± SEM. The fitted parameters including **(B)** normalized amplitude, **(C)** peak frequency and **(D)** bandwidth, were compared between groups. Error bars represent 95% confidence intervals. ***$P < 0.01$* (*F*-test).

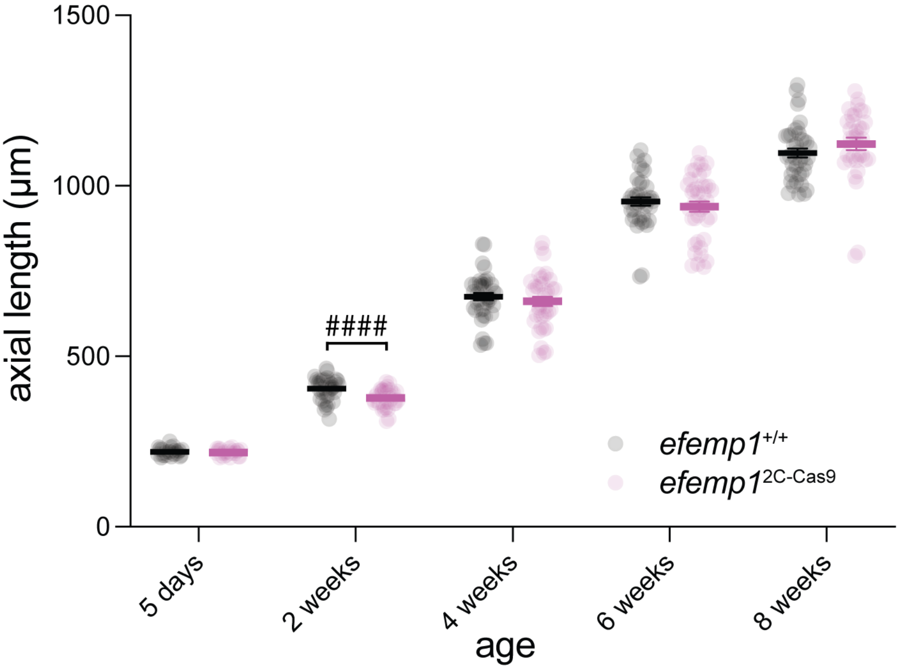


Figure S2. **Axial length of *efemp1*$^{+/+}$ and *efemp1*$^{2C\text{-}Cas9}$ fish under normal lighting.** Axial length of *efemp1*$^{+/+}$ and *efemp1*$^{2C\text{-}Cas9}$ fish at 5 days post-fertilization (dpf; n = 30 and 29 eyes, respectively), and 2 (n = 40 per genotype), 4 (n = 40 and 38 eyes, respectively), 6 (n = 39 and 40 eyes, respectively), and 8 weeks post-fertilization (wpf; n = 40 and 34 eyes, respectively). *Two*-way ANOVA was performed. Group data are shown as mean ± SEM. *Two*-way ANOVA, Fisher's LSD *post-hoc* tests (*$P < 0.05$; **$P < 0.01$; ***$P < 0.001$; ****$P < 0.0001$) and multiple unpaired *t*-test ($^{####}P < 0.0001$) were conducted. n.s.: not significant.

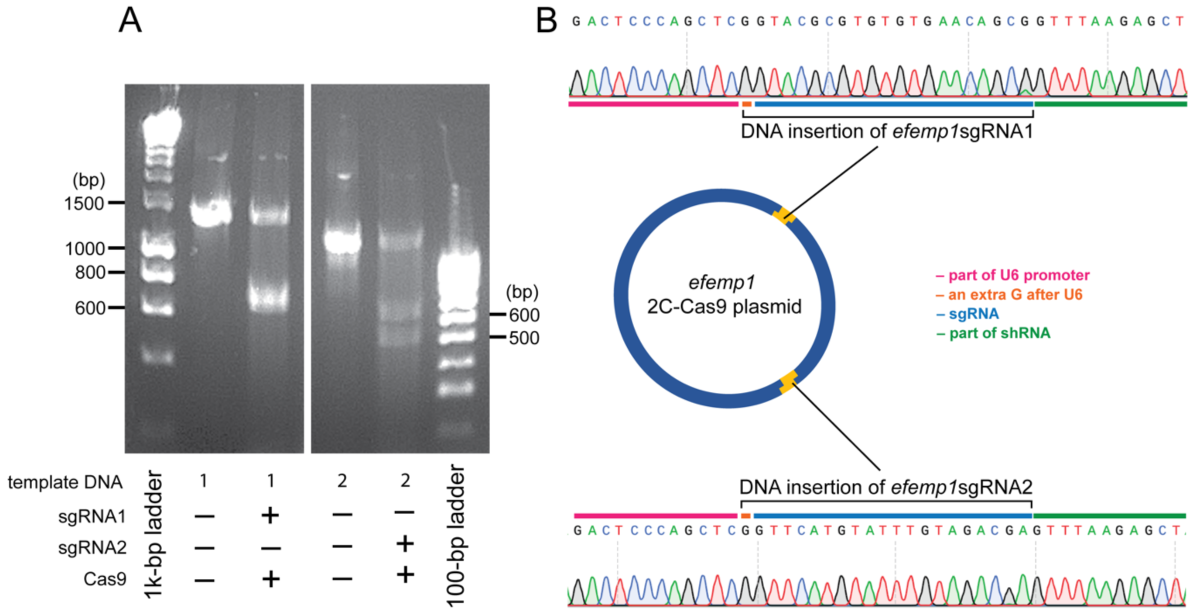


**Figure S3. *In vitro* assessment of *efemp1* sgRNAs (A) and confirmation of DNA insertions of *efemp1* sgRNAs in plasmid (B).** **(A)** SgRNAs targeting *efemp1* gDNA for CRISPR gene editing was tested. On the left gel image, from left to right, the first lane presents 1-kb marker ladder. The second lane shows purified PCR products (~120 ng) of a DNA sequence (1373 base pairs, bp) containing the target sites of *efemp1* sgRNA1. In the third lane, Cas9 nuclease (50 ng) binds with *efemp1* sgRNA1 (75 ng) to cut the DNA sequence to 712 and 661 bp bands (the two bands overlay on the gel). Positions of 600, 800, 1000 and 1500 bp were highlighted on the left of the gel image. On the right gel image, from left to right, the first lane shows purified PCR products (~120 ng) of a DNA sequence (1141 bp) containing the target sites of *efemp1* sgRNA2. In the second lane, Cas9 nuclease (50 ng) binds with *efemp1* sgRNA2 (75 ng) to cut the DNA sequence to 507 and 634 bp bands. The third lane shows a 100-bp marker ladder. Positions of 500 and 600 bp were highlighted on the right of the gel image. **(B)** Sanger sequencing results confirms the insertions of sgRNA DNA sequences (highlighted in blue lines) into the 2C-Cas9 plasmid.

Table S1. **Proportion of *efemp1*$^{2C\text{-}Cas9}$ fish with enlarged eyes**

| Tank | Date of Birth | Checking date | Number of fish with enlarged eyes | Fish number of the tank | % of fish with enlarged eyes | Average (%) |
|---|---|---|---|---|---|---|
| 1 | 05.02.2022 | 10.10.2022 | 4 | 28 | 14.2857143 | |
| 2 | 05.02.2022 | 10.10.2022 | 0 | 30 | 0 | |
| 3 | 11.03.2022 | 10.10.2022 | 5 | 35 | 14.2857143 | 7.10963455 |
| 4 | 06.05.2022 | 10.10.2022 | 3 | 43 | 6.97674419 | |
| 5 | 21.04.2022 | 10.10.2022 | 0 | 18 | 0 | |

Table S2. **Oligos to re-anneal to the double-strand DNA sequences of *efemp1* sgRNAs to insert into the 2C-Cas9 plasmid**

| SgRNA | Forward (5′ to 3′) | Reverse (5′ to 3′) |
|---|---|---|
| Efemp1sgRNA1 | CTCG**GTACGCGTGTGTGAACAGCG** | AAAC**CGCTGTTCACACACGCGTAC** |
| Efemp1sgRNA2 | AGCTCG**GTTCATGTATTTGTAGACGA**G | TAAAC**TCGTCTACAAATACATGAAC**CG |

Note: Efemp1sgRNA1 was inserted into the site digested by BsaI enzyme. Efemp1sgRNA2 was inserted into the site digested by BsmBI enzyme. Bold fonts highlight sgRNA sequences and regular fonts form sticky ends to specifically bind to digested sites of the plasmid.

Table S3. **Primers for genotyping *efemp1*$^{2C\text{-}Cas9}$ fish**

| Target site | PCR | Forward (5′ to 3′) | Reverse (5′ to 3′) |
|---|---|---|---|
| 1 | standard | CTTCAGGTCCATTTGACATCC | AGCTCCAGTGGCAGGGTT |
| 1 | headloop | CTTCAGGTCCATTTGACATCC | **TGTGAACAGCGAGGGTGGATACT**AGCTCCAGTGGCAGGGTT |
| 2 | standard | TCTTTGTGCTTCCTCCTTTG | TCCGTCATTGCCGTCTTT |
| 2 | headloop | TCTTTGTGCTTCCTCCTTTG | **TCCATCGTCTACAAATACAT**TCCGTCATTGCCGTCTTT |

Note: Bold fonts indicate the extension (headloop tag) of the primer for headloop PCR.
